# Distinct proteomic and acylproteomic adaptations to succinate dehydrogenase loss in two cell contexts

**DOI:** 10.1101/2025.08.01.668168

**Authors:** Sherry X Zhou, Benjamin Madden, M Cristine Charlesworth, Taro Hitosugi, Judith Favier, L. James Maher

## Abstract

**Background:** The tricarboxylic acid (TCA) cycle and electron transport chain (ETC) are key metabolic pathways required for cellular ATP production. While loss of components in these pathways typically impairs cell survival, such defects can paradoxically promote tumorigenesis in certain cell types. One such example is loss of succinate dehydrogenase (SDH), which functions in both the TCA cycle and as Complex II of the ETC. Deleterious mutations in SDH subunits can cause pheochromocytoma and paraganglioma (PPGL), rare hereditary neuroendocrine tumors of chromaffin cells in the adrenal gland and the nerve ganglia, respectively. Why tumor formation upon SDH loss is limited to certain tissues remains unclear. We hypothesized that the metabolic and proteomic perturbations resulting from SDH loss are cell-type specific, favoring survival of chromaffin cells.

**Methods:** We comprehensively examined the proteomic, acetylproteomic, and succinylproteomic effects of SDH loss in two cell models, immortalized mouse chromaffin cells (imCCs) and immortalized mouse embryonic fibroblasts (iMEFs). Perturbations in metabolite levels were determined by mass spectrometry. Effects of SDH loss on fatty acid β-oxidation (FAO) were assessed by stable isotope tracing and pharmacologic inhibition.

**Results:** SDH-loss imCCs show significant upregulation of mitochondrial proteins, including TCA cycle and FAO enzymes, with pronounced downregulation of nuclear proteins. Both imCCs and iMEFs demonstrate significant energy deficiency upon SDH loss, but FAO activity is uniquely increased in SDH-loss imCCs. While SDH loss increases both lysine-reactive acetyl-CoA and succinyl-CoA, SDH-loss imCCs and iMEFs show disproportionate hyperacetylation but mixed succinylation. Surprisingly, SDH-loss imCCs, but not iMEFs, display disproportionate hypoacetylation and hyposuccinylation of mitochondrial proteins.

**Conclusions:** SDH loss differentially impacts the proteomes and acylproteomes of imCCs and iMEFs, with compartment-specific effects. These findings reveal cell type-specific adaptations to SDH loss. The plasticity of the response of imCCs may underlie the tissue-specific susceptibility to tumorigenesis and could illuminate therapeutic vulnerabilities of SDH-loss tumors.

## Background

The tricarboxylic acid (TCA) cycle and the electron transport chain (ETC) are key pathways that maintain cellular energy and metabolite homeostasis. Inputs from glycolysis and lipid catabolism feed into the TCA cycle, supporting ATP production via the ETC and providing precursors for metabolite and fatty acid synthesis. The TCA cycle and ETC are connected in two main ways: by the electron carrier NADH via Complex I, and directly through the heterotetrameric succinate dehydrogenase (SDH) enzyme complex, which functions in both the TCA cycle and as Complex II of the ETC.

Despite the critical role of the TCA cycle in maintaining energy and metabolite pools, loss of TCA cycle enzymes can, paradoxically, promote tumorigenesis of gliomas, neuroendocrine tumors, and renal cell carcinoma. Deleterious mutations in SDH subunits and its assembly factors, as well as *MDH2*, *FH*, *IDH3B*, and *DLST*, predispose individuals to pheochromocytoma and paraganglioma (PPGL), rare neuroendocrine tumors of chromaffin cells in the adrenal gland and the nerve ganglia, respectively (1), indicating a unique link between TCA cycle dysfunction and PPGL tumorigenesis. However, the tissue specificity of SDH-loss driven tumors, particularly SDH-loss PPGL, which is the second most common inherited cause of PPGL (2), remains poorly understood.

The SDH complex is composed of four nuclear-encoded subunits (SDHA-D) and requires four assembly factors (SDHAF1-4) for proper maturation (3, 4). Mutations disrupting any subunit render the complex inactive. Carriers heterozygous for deleterious SDH subunit variants are unaffected but are susceptible to PPGL tumor initiation upon somatic loss of the remaining functioning allele.

A leading model for PPGL tumorigenesis driven by SDH deficiency posits that TCA cycle disruption leads to accumulation of intermediates that act as oncometabolites. In particular, succinate buildup following SDH loss can competitively inhibit dozens of 2-ketoglutarate-dependent dioxygenases that produce succinate as a byproduct (5–8). Among these vulnerable dioxygenases are Jumonji domain histone demethylases and TET-family DNA cytosine demethylases, whose inhibition results in histone and DNA hypermethylation (8–11). Other vulnerable enzymes include prolyl hydroxylases required for collagen maturation and for marking hypoxia-inducible transcription factors (HIFs) for oxygen-dependent degradation (5, 6, 8, 11). Impaired proteolysis of HIFs in the presence of oxygen drives a “pseudohypoxic” phenotype. Notably, this pseudohypoxia is observed in both SDH-loss and VHL-loss PPGLs, the latter unable to degrade HIFs due to a downstream defect in proteolysis. Interestingly, SDH-loss PPGLs have been found to be more aggressive than VHL-loss tumors (12), suggesting that SDH loss promotes tumor growth through additional mechanisms beyond pseudohypoxia. These may include the other succinate toxicities mentioned above as well as increased production of reactive oxygen species (13–17).

We have been pursuing two main questions: (i) whether additional pathogenic mechanisms beyond succinate poisoning of dioxygenase enzymes explain the increased pathogenicity of SDH-loss PPGLs, and (ii) what serves as the basis for the unique tissue specificity of PPGL tumorigenesis.

In addition to succinate accumulation, TCA cycle blockade due to SDH loss may lead to the accumulation of other upstream metabolites. Acetyl-CoA and succinyl-CoA came to our attention as highly reactive metabolites capable of post-translationally acylating the lysine side chain ε-amino group in both enzymatic and spontaneous chemical reactions, resulting in acetyllysine and succinyllysine, respectively (18–21). Histone lysine acylation is a well-studied example of protein acylation with epigenetic consequences (18, 19, 22). Acylation of non-histone proteins has also been shown to affect protein stability and function (23–30). Protein hypersuccinylation upon SDH loss is plausible: SDH inhibition, knockdown, or deletion, or introduction of exogenous cell-permeable succinate or 2-ketoglutarate analogs have all been shown to increase succinyl-CoA levels in various cell models (22, 30, 31). Similarly, loss of the TCA cycle enzyme succinyl-CoA synthetase raises succinyl-CoA levels in cell models and patient samples (32). Importantly, multiple studies have demonstrated the association of protein acylation with disease states, suggesting that accumulation of succinyl-CoA and acetyl-CoA predispose cells to pathological hyperacylation (22, 31, 33). However, broader implications of altered protein acylation in the context of metabolic dysregulation remain poorly understood, and specific effects in PPGL are yet to be explored.

Here, we performed TMT-labeled quantitative proteomic and acylproteomic analyses to comprehensively examine the effects of SDH loss in mouse adrenal-derived cells compared to fibroblasts. We sought to determine how SDH loss (i) alters the proteome and (ii) alters lysine acetylation and succinylation across the proteome (the acetylome and succinylome, respectively). We exploited available models, namely SDH-containing vs. SDHC-loss immortalized mouse fibroblasts (iMEFs) (34) and SDH-containing vs. SDHB-loss immortalized mouse adrenal cells (imCCs) (35). We note that the ideal comparison between cell types would involve loss of the same SDH subunit, but such paired models are not available.

We set out to test two hypotheses. First, we hypothesized that the tissue preference for PPGL tumorigenesis in adrenal-derived cells rather than fibroblasts would manifest as a distinct proteomic response to SDH loss in imCCs, reflecting a greater capacity for adrenal cells to survive the metabolic insult (36, 37). Indeed, our results support this hypothesis. We find that the imCC proteome undergoes more extensive alterations upon SDH loss than the iMEF proteome. Shared patterns include upregulation of collagen biosynthesis and downregulation of translation-related pathways and cell cycle regulation. We observe compartment-specific changes upon SDH loss, with imCCs significantly upregulating mitochondrial proteins and downregulating nuclear proteins. Notably, TCA cycle and fatty-acid β-oxidation (FAO) enzymes are upregulated in SDH-loss imCCs, suggesting that these cells possess unique metabolic and proteomic plasticities, including the activation of FAO, that may enhance their capacity to generate energy for survival. Second, we hypothesized that lysine-reactive acetyl-CoA and succinyl-CoA are increased in SDH-loss cells, leading to global protein hyperacylation. Surprisingly, while accumulation of these acyl-CoAs was indeed confirmed upon SDH loss, alterations in protein acetylation and succinylation were mixed, with more pronounced alterations in imCCs upon SDH loss. We find that disproportionate protein acylation changes vary strikingly across subcellular compartments, revealing much more complex regulation of post-translational lysine acylation than hypothesized.

Together, our findings suggest that cell-type specific proteomic and acyl-proteomic adaptations to SDH loss favor survival of specific cell types (here, adrenal-derived cells) that can better compensate for defective energy metabolism. This observation may help explain the selective vulnerability of certain tissues to SDH-loss-driven tumorigenesis.

## Results

### SDH loss differentially alters the proteomes of imCCs and iMEFs

To understand the effects of SDH loss on the proteomes of imCCs and iMEFs, we performed TMT-based proteomic analyses of imCCs and iMEFs with and without SDH loss (Supplemental Fig. S1). Principal component analysis confirmed that results group by SDH-loss status within each cell type, as expected (Supplemental Fig. S2A). Interestingly, SDH loss had a greater impact on the imCC proteome, with 34% of proteins significantly altered [fold change ≥ 2; false-discovery rate (FDR) adjusted p-value < 0.05] as opposed to 18% in iMEFs (Fig. 1A). Of the 8187 proteins detected in imCCs and 7925 proteins detected in iMEFs, 7322 proteins were detected in both (Fig. 1B). Notably, only about one quarter of these shared proteins were comparably affected by SDH loss. A large fraction of shared proteins (41%) displayed greater than 1.25-fold differential increase in imCCs vs. iMEFs, while 34% of shared proteins displayed greater than 1.25-fold differential increase in iMEFs vs. imCCs (Fig. 1C). As expected, a comparison with our previous lower-resolution SILAC proteomics analysis of control and SDH-loss iMEFs (22) showed that over 98% of proteins previously identified were also detected in the present iMEF analysis, and measured fold changes are positively correlated between the two different studies (Supplemental Fig. S2B).

**Figure 1.**
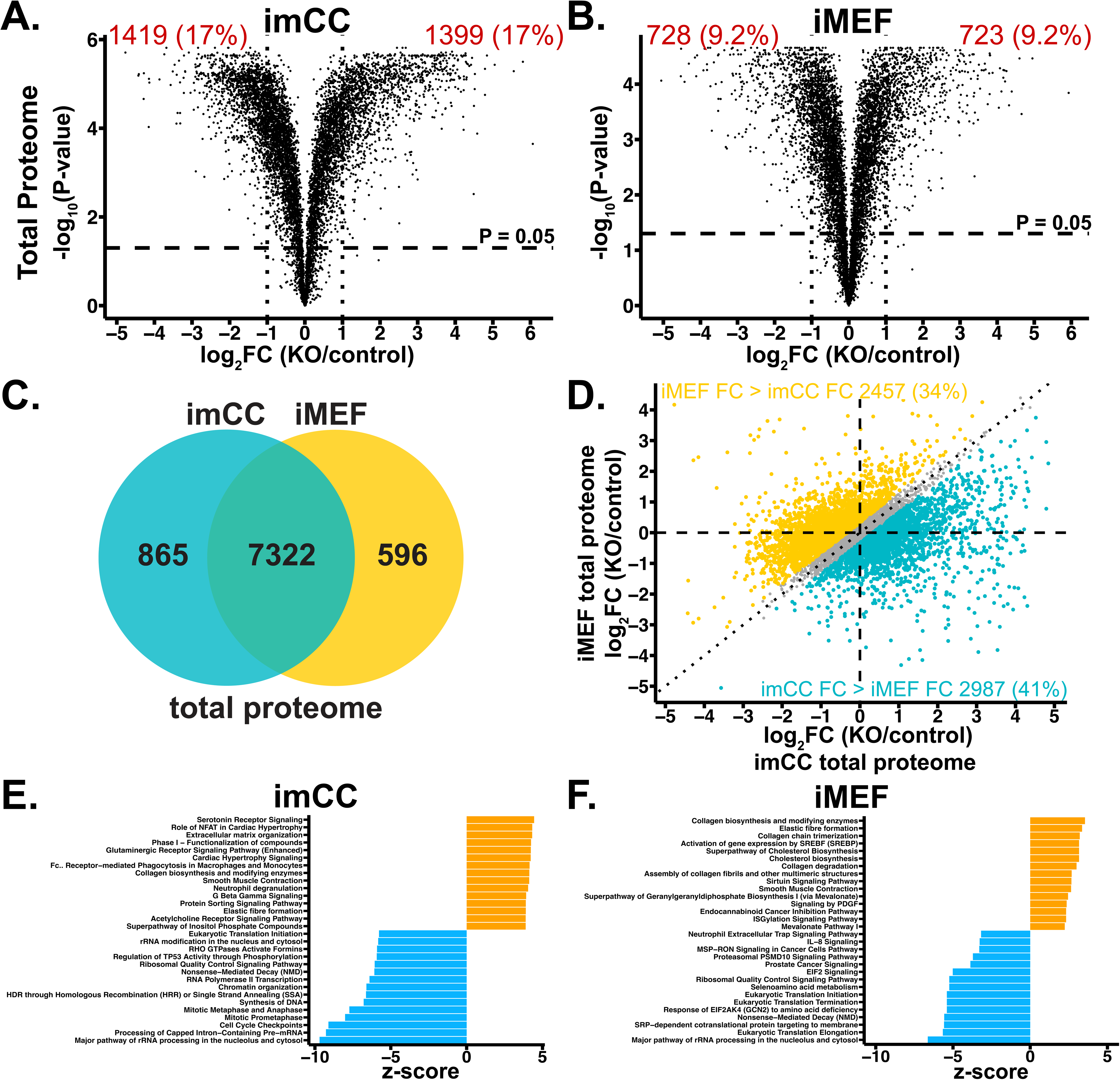
SDH loss significantly alters imCC and iMEF total proteomes. **A-C.** Volcano plot depicting effects of SDH loss on **A.** imCC and **B.** iMEF total proteomes. Vertical dotted lines indicate fold change (FC) of 2. The number and percentage of significantly changed (P (FDR-adjusted p-value) < 0.05, |FC| > 2) proteins are labeled in red. **C.** Venn diagram of all identified proteins shown in **A.** and **B.** as shared or unique to imCC and iMEF total proteomes. **D.** Correlation plot of total protein changes upon SDH loss between imCCs and iMEFs. Proteins with iMEF FC/imCC FC > 1.25 are colored in gold and proteins with imCC FC/iMEF FC > 1.25 are colored in teal. **E-F.** Top 15 upregulated and downregulated pathways upon SDH loss for **E.** imCCs and **F.** iMEFs from Ingenuity Pathway Analysis.

Ingenuity pathway analysis (IPA) (38) of proteins whose levels were significantly altered upon SDH loss showed both shared and distinct pathway changes in the functional proteomes of imCCs and iMEFs. Shared changes in the top 15 dysregulated pathways for both cell types include upregulation of collagen biosynthesis and downregulation of translation, rRNA processing, and cell cycle control (Fig. 1D, E). Upregulation of collagen biosynthesis upon SDH loss is consistent with our previous finding of decreased trypsin sensitivity in SDH-loss imCCs and iMEFs (37). Interestingly, these top dysregulated protein pathways differ from those identified by transcriptome analysis (37), indicating that mRNA levels are not directly reflective of protein expression in these cell models. Indeed, this discrepancy may be related to the observed profound downregulation of translation and rRNA processing-related pathways upon SDH loss. We find that in the 7923 imCC and 7658 iMEF mRNA-protein pairs identified (Supplemental Fig. S3A-B), changes induced upon SDH loss show overall positive correlations. However, interestingly, quantitative effects of SDH loss are greater in the proteome than in the transcriptome (Supplemental Fig. S3C-D).

Among the 42 altered pathways shared between SDH-loss imCCs and iMEFs (Supplemental Fig. S4A), only one dysregulated pathway, regulation of lipid metabolism by PPARα, was downregulated in imCCs while upregulated in iMEFs (Supplemental Fig. S4B). In imCCs, over half of the dysregulated proteins associated with this pathway are Mediator complex subunits, suggesting that SDH-loss imCCs, but not iMEFs, may have decreased transcription in addition to decreased translation. Upon SDH loss, sixteen pathways were upregulated in imCCs and downregulated in iMEFs. Most such pathways were related to angiogenesis and cancer cell signaling (Supplemental Table S1). Indeed, while a pseudohypoxic phenotype is expected in both imCCs and iMEFs due to succinate accumulation and stabilization of HIF proteins by inhibition of prolyl hydroxylation required for degradation (5, 6, 8), hypoxic response pathways appear to be more activated by SDH loss in imCCs.

IPA for SDH loss effects also revealed diverging phenotypes related to mitochondrial function (Supplemental Fig. S4C). Five mitochondrial pathways (mitochondrial fatty acid β-oxidation, TCA cycle enzyme maturation/regulation, mitochondrial protein degradation, mitochondrial iron-sulfur cluster biogenesis, mitochondrial division signaling) were upregulated in imCCs upon SDH loss. In stark contrast, no mitochondrial pathways were significantly dysregulated in SDH-loss iMEFs, demonstrating that SDH loss leads to much more dramatic mitochondrial proteomic remodeling in imCCs compared to iMEFs. Together, these data suggest that SDH loss significantly alters the proteomes of both imCCs and iMEFs, leading to shared downregulation of cell cycle regulation and protein translation and a unique upregulation of mitochondrial proteins in imCCs.

### SDH-loss imCCs display subcellular compartment-specific proteomic changes

We further explored subcellular compartment-specific alterations induced by SDH loss by analyzing levels of proteins localized to the mitochondria, nucleus, and cytoplasm. Notably, only imCCs show a significant upregulation of mitochondrial proteins upon SDH loss (Fig. 2A). Moreover, downregulation of nuclear protein expression was more prominent upon SDH loss in imCCs than in iMEFs (Fig. 2B). DAVID functional annotation of nuclear proteins significantly downregulated upon SDH loss in imCCs shows particular enrichment of pathways related to DNA damage and repair, mRNA and rRNA processing, and DNA replication and cell division relative to iMEFs (Supplemental Fig. S5A-B). Interestingly, succinate is an oncometabolite reported to inhibit one or more key enzymes in DNA repair (39), suggesting that its accumulation upon SDH loss might be related to downregulation of DNA repair machinery. Downregulation of DNA replication and cell division pathways upon SDH loss is consistent with the markedly increased doubling times of SDH-loss imCCs and iMEFs (37).

**Figure 2.**
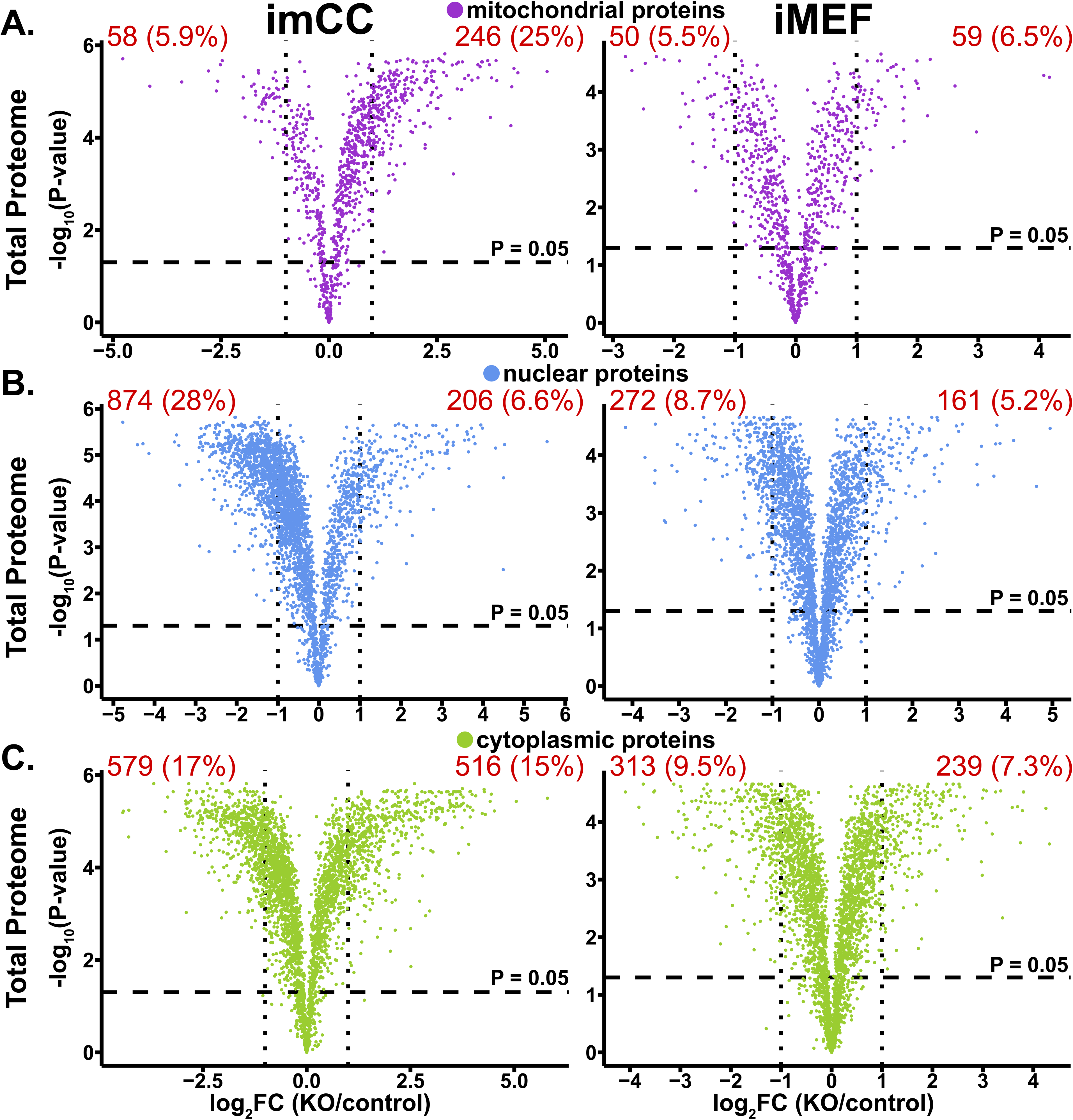
SDH loss leads to greater compartment-specific proteome changes in imCCs than in iMEFs. **A-C.** Volcano plots depicting SDH-loss-induced proteome changes in **A.** mitochondrial, **B.** nuclear, and **C.** cytoplasmic proteins in imCCs and iMEFs. Vertical dotted lines indicate FC of 2. The number and percentage of significantly changed (P (FDR-adjusted p-value) < 0.05, |FC| > 2) proteins are labeled in red.

In contrast to the distinct changes for mitochondrial and nuclear proteins in SDH-loss imCCs vs. iMEFs, similar numbers of cytoplasmic proteins were significantly increased or decreased upon SDH loss in imCCs and iMEFs (Fig. 2C). These compartment-specific variations in proteomic responses to SDH loss suggest that different cell types respond uniquely to SDH loss and may rely upon distinct compensatory metabolic pathways.

### SDH loss induces a profound low-energy cellular state

The striking slowing of cell division and downregulation of translation-related processes upon SDH loss suggest that this TCA cycle dysfunction induces uncompensated energy deficiency. We confirmed this hypothesis by measuring cellular energy balance. Both SDH-loss imCCs and iMEFs demonstrated profoundly decreased ATP/ADP and increased NAD^+^/NADH ratios, indicative of low metabolic energy currency and an oxidizing redox status upon SDH loss (Fig. 3A-D). Notably, the low ATP/ADP ratio in imCCs is primarily due to a three-fold increase in ADP concentration upon SDH loss while in iMEFs the altered ratio is primarily driven by a four-fold decrease in ATP upon SDH loss. Similarly, the high NAD^+^/NADH ratio in SDH-loss imCCs is driven by a 1.75-fold increase in NAD^+^, while in SDH-loss iMEFs is driven by a three-fold decrease in NADH. Thus, although both cell types suffer severe energy depletion upon SDH loss, the different origins of energy carrier imbalances between imCCs and iMEFs suggest cell-type specific adaptations to energy stress induced by this TCA cycle defect.

**Figure 3.**
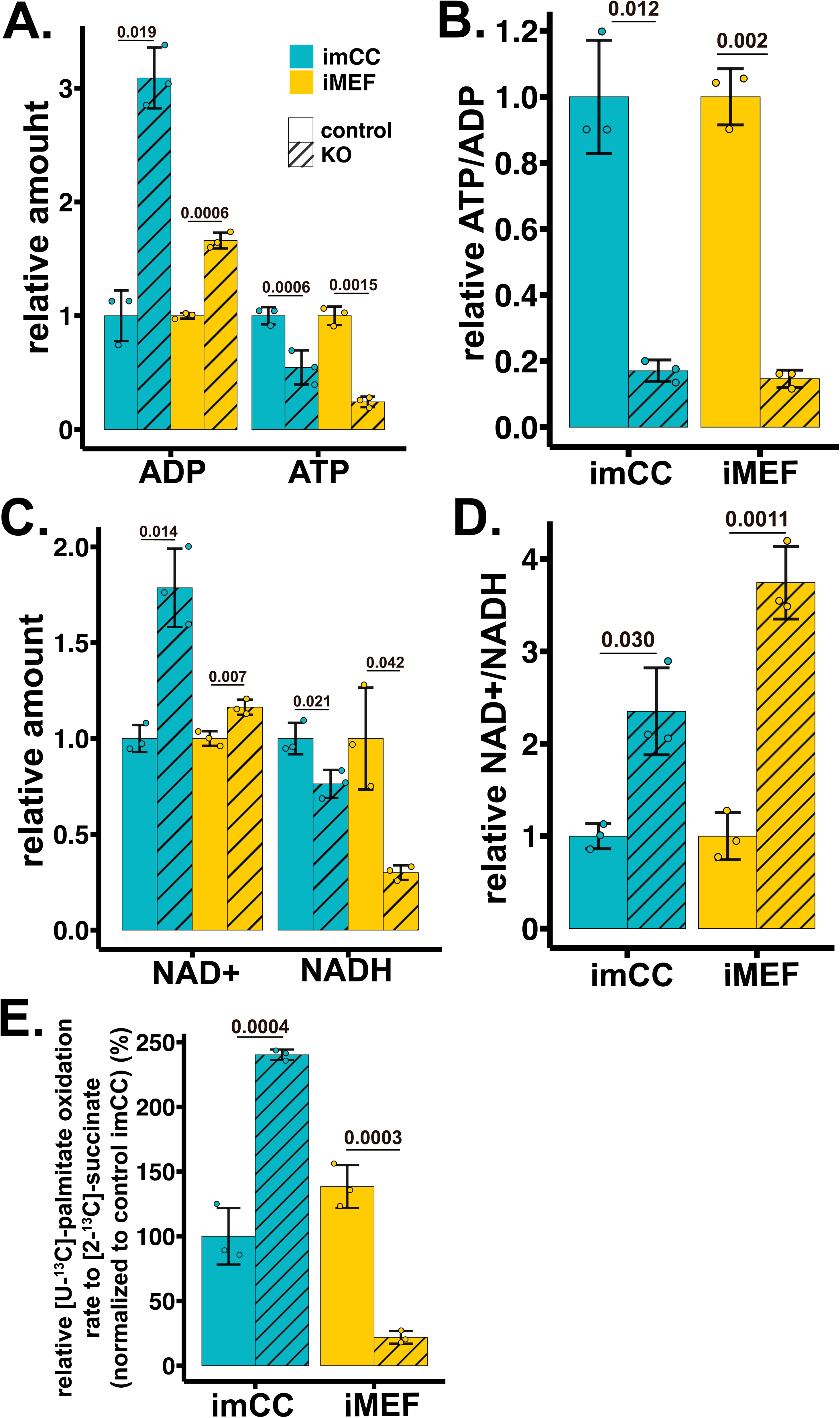
SDH loss leads to significant energy deficiency in imCCs and in iMEFs. **A.** ATP and ADP in imCCs and iMEFs measured by liquid chromatography-mass spectrometry (LCMS) shown as amount relative to controls. **B.** ATP/ADP ratio in imCCs and iMEFs relative to controls. **C.** NAD^+^ and NADH in imCCs and iMEFs measured by LCMS shown as amount relative to controls. **E.** NAD^+^/NADH ratio in imCCs and in iMEFs relative to controls. **F.** Relative [M+2]-labeled [^13^C] succinate oxidation from [U-^13^C]-palmitate.

### Cell-type specific alterations occur in the mitochondrial proteome upon SDH loss

To determine what mitochondrial pathways were most affected in SDH-loss imCCs, we performed DAVID functional annotation (40, 41) of significantly upregulated mitochondrial proteins. Notably, fatty acid metabolism-related pathways were the top significantly enriched clusters (with much higher statistical significance for SDH-loss imCCs than iMEFs). TCA cycle and FAD-binding proteins were also significantly enriched (Fig. 4A, B). Independent mass spectrometry quantification of FAD, an electron carrier essential for fatty acid oxidation (FAO), showed a greater than two-fold increase unique to SDH-loss imCCs (Fig 4C, D). Upon SDH loss, FAD-binding FAO-related proteins were also upregulated in imCCs compared to iMEFs, suggesting that higher FAD levels may support increased FAO in SDH-loss imCCs (Fig. 4E, F). We also found that SDH loss triggered a greater increase of mitochondrial FAO proteins in imCCs than in iMEFs (Fig. 4G, H).

**Figure 4.**
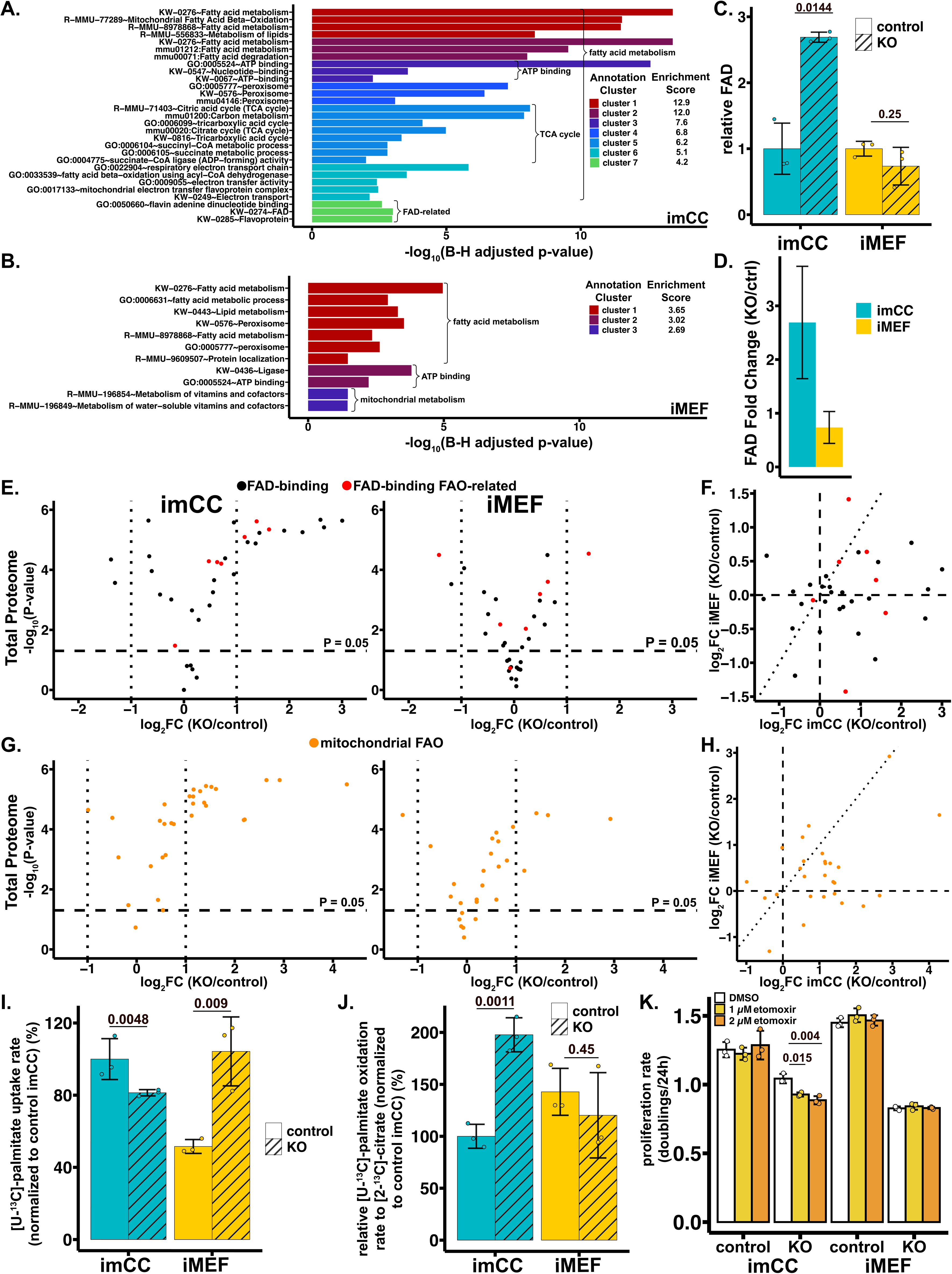
SDH-loss imCCs upregulate fatty acid β-oxidation and FAD-binding proteins more than SDH-loss iMEFs. **A-B.** Top annotation clusters from DAVID Functional Analysis of mitochondrial proteins with FC > 2, P < 0.05 in **A.** imCCs and **B.** iMEFs. **C.** FAD in imCCs and iMEFs measured by LCMS (mean ± SD, *n* = 3 biological replicates per condition). P-values were calculated using two-tailed Welch’s t-tests. **D.** Fold change of FAD levels in **C.** upon SDH loss**. E.** Volcano plots depicting SDH-loss-induced changes in FAD-binding proteins in total proteomes of imCCs and iMEFs. FAD-binding proteins shown in black; FAD-binding FAO-related proteins shown in red. **F.** Correlation plot comparing SDH-loss-induced changes in FAD-binding proteins between imCCs and iMEFs. **G.** Volcano plots depicting SDH-loss-induced changes in fatty acid β-oxidation proteins (Reactome R-MMU-77289) in total proteomes of imCCs and iMEFs. **H.** Correlation plot comparing SDH-loss-induced changes in fatty acid β-oxidation proteins between imCCs and iMEFs. **I.** [U-^13^C]-palmitate uptake rate normalized to control imCC (mean ± SD, *n* = 3 biological replicates per condition). P-values were calculated using two-tailed Welch’s t-tests. **J.** FAO activity measured by [U-^13^C]-palmitate oxidation rate to [M+2]-labeled [^13^C] citrate (mean ± SD, *n* = 3 biological replicates per condition). P-values were calculated using two-tailed Welch’s t-tests. **K.** Proliferation rates of vehicle (DMSO) and etomoxir-treated imCCs and iMEFs in modified culture media with reduced glucose (5 mM) and reduced glutamine (0.55 mM). Cells were treated at indicated concentrations for 3 cell doublings. P-values were calculated using two-tailed Welch’s t-tests. Data representative of 3 independent replicates.

To assess if imCCs, but not iMEFs, have functionally increased FAO activity upon SDH loss, we monitored conversion of isotopically labeled palmitate to citrate. Intriguingly, SDH loss resulted in significantly lower palmitate uptake in imCCs and significantly higher uptake in iMEFs (Fig. 4I). On the other hand, SDH-loss imCCs exhibited almost twice the normalized conversion rate of labeled palmitate to citrate compared to control, while there was no significant change in conversion rate in iMEFs upon SDH loss (Fig. 4J). These results suggest that SDH-loss imCCs take up fatty acids for energy production through FAO while SDH-loss iMEFs take up fatty acids for other processes such as membrane biosynthesis. We therefore hypothesized that growth of SDH-loss imCCs would be more sensitive to FAO inhibition than growth of SDH-loss iMEFs, particularly since the key FAO regulator CPT1A is five-fold higher in imCCs upon SDH loss while not significantly changed upon SDH loss in iMEFs (Supplemental Fig. 6A). We treated control and SDH-loss imCCs and iMEFs for three doublings with etomoxir, an irreversible inhibitor of CPT1A. Etomoxir treatment selectively decreased the proliferation rate of SDH-loss imCCs, with a more pronounced effect under more physiological conditions (Fig. 4K, Supplemental Fig. S6B), suggesting that SDH-loss imCCs indeed display a measurable dependence on FAO for maintaining cellular proliferation.

We take this increased dependence of SDH-loss imCCs on upregulation of FAO for proliferation as evidence that SDH-loss imCCs derive energy from FAO-derived NADH and FADH_2_, which typically enter the ETC through Complex I (NADH) and Complex III via ETFDH (for acyl-CoA dehydrogenase-bound FADH_2_). Interestingly, despite previous transcriptomic findings showing upregulation of a subset of Complex I genes in imCCs but not iMEFs upon SDH loss, and preservation of Complex I activity in SDH-loss imCCs but not iMEFs (37), our proteomic data reveal that most Complex I subunits are downregulated in both imCCs and iMEFs, suggesting that Complex I activity is not solely dependent on protein abundance (Supplemental Fig. S7A). In contrast, most Complex III subunits are significantly decreased upon SDH loss in iMEFs but not in imCCs, some Complex IV subunits are increased in iMEFs but not in imCCs, and Complex V subunits are significantly upregulated in imCCs while downregulated in iMEFs (Supplemental Fig. S7B-E). Given that Complex III activity is required for recycling of the coenzyme Q pool, which carries electrons not only from Complex I and II but also ETFDH, which receives electrons from FAO, branched chain amino acid (BCAA) catabolism, and sarcosine metabolism, its loss may impair ETC function and ATP production in SDH-loss iMEFs. Complex III has also been implicated as important for tumor growth (42), suggesting that SDH-loss imCCs may be more metabolically fit than SDH-loss iMEFs. Thus, even in the face of TCA cycle dysfunction and aberrant mitochondria, Complex V upregulation may facilitate some ATP generation in SDH-loss imCCs. This finding suggests that, despite their low-energy status, these cells may benefit from greater metabolic plasticity.

In addition to the upregulation of FAO, SDH-loss imCCs, but not iMEFs, upregulate multiple TCA cycle proteins in response to SDH loss (Supplemental Fig. S7F). Although some TCA cycle metabolite levels were significantly reduced upon SDH loss in both cell types (37), SDH-loss imCCs appear to retain the ability to generate some energy from TCA cycle steps upstream of SDH. In particular, the conversion of succinyl-CoA to succinate by succinyl-CoA ligase produces ATP or GTP, and we found that levels of succinyl-CoA ligase subunits were elevated in imCCs but not iMEFs upon SDH loss (Supplemental Fig. S7F). We also found that imCCs produce more labeled succinate from labeled palmitate upon SDH loss, while iMEFs produce much less labeled succinate despite similar production of labeled citrate upon SDH loss (Fig. 4E). These results suggest that SDH-loss iMEFs divert citrate away from the TCA cycle while imCCs do not, and that SDH-loss imCCs again display enhanced metabolic plasticity in their ability to generate additional energy via the portion of the TCA cycle upstream of SDH.

### SDH loss leads to mixed alterations in protein acetylation and succinylation

Loss of SDH has been shown to alter the levels of various TCA cycle metabolites upstream and downstream of SDH (22, 37, 43–45). However, the levels of two reactive TCA cycle metabolites, acetyl-CoA and succinyl-CoA, have not been routinely quantified. Acetyl-CoA and succinyl-CoA can both either spontaneously or enzymatically react with lysine side chains, altering lysine charge by acetylation and succinylation (Fig. 5A). These metabolites directly connect cellular metabolic state to proteome acylation status, with potential effects on protein function and epigenetic regulation. We therefore performed quantitative mass spectrometry to assess the impact of SDH loss on free acetyl-CoA and succinyl-CoA levels in imCCs and iMEFs. Both cell types showed significantly increased acetyl-CoA and succinyl-CoA levels upon SDH loss, as we originally hypothesized, (Fig. 5B), indicating that succinate is not the only TCA cycle metabolite that accumulates upon SDH loss. Because acetyl-CoA and succinyl-CoA are required substrates for lysine acylation, we originally predicted global protein hyperacetylation and hypersuccinylation in SDH-loss cells. However, we then considered that post-translational acetylation and succinylation marks can be reversed enzymatically by deacylating enzymes such as sirtuins and histone deacetylases (HDACs), many of which require NAD^+^ as a cofactor (Fig. 5A). We therefore independently quantified NAD^+^ by mass spectrometry and showed, notably, that NAD^+^ levels were two-fold higher in imCCs and slightly increased in iMEFs upon SDH loss (Fig. 3C). This result raised the possibility that SDH-loss protein acylation might be balanced to some extent by NAD^+^-driven enzymatic (sirtuin and HDAC) deacylation. To directly assess sirtuin activity in cultured imCCs and iMEFs, we implemented a live-cell dual mCherry-EGFP(K85AcK) reporter system in which deacetylation results in EGFP fluorescence (46). Indeed, flow cytometry quantification of imCCs carrying the fluorescent reporters showed significantly increased sirtuin deacetylase activity upon SDH loss, while SDH-loss iMEFs show significantly decreased sirtuin deacetylase activity (Fig. 5C). These results suggest that elevated NAD^+^ in SDH-loss imCCs promotes increased sirtuin activity, with the potential to mitigate protein hyperacylation.

**Figure 5.**
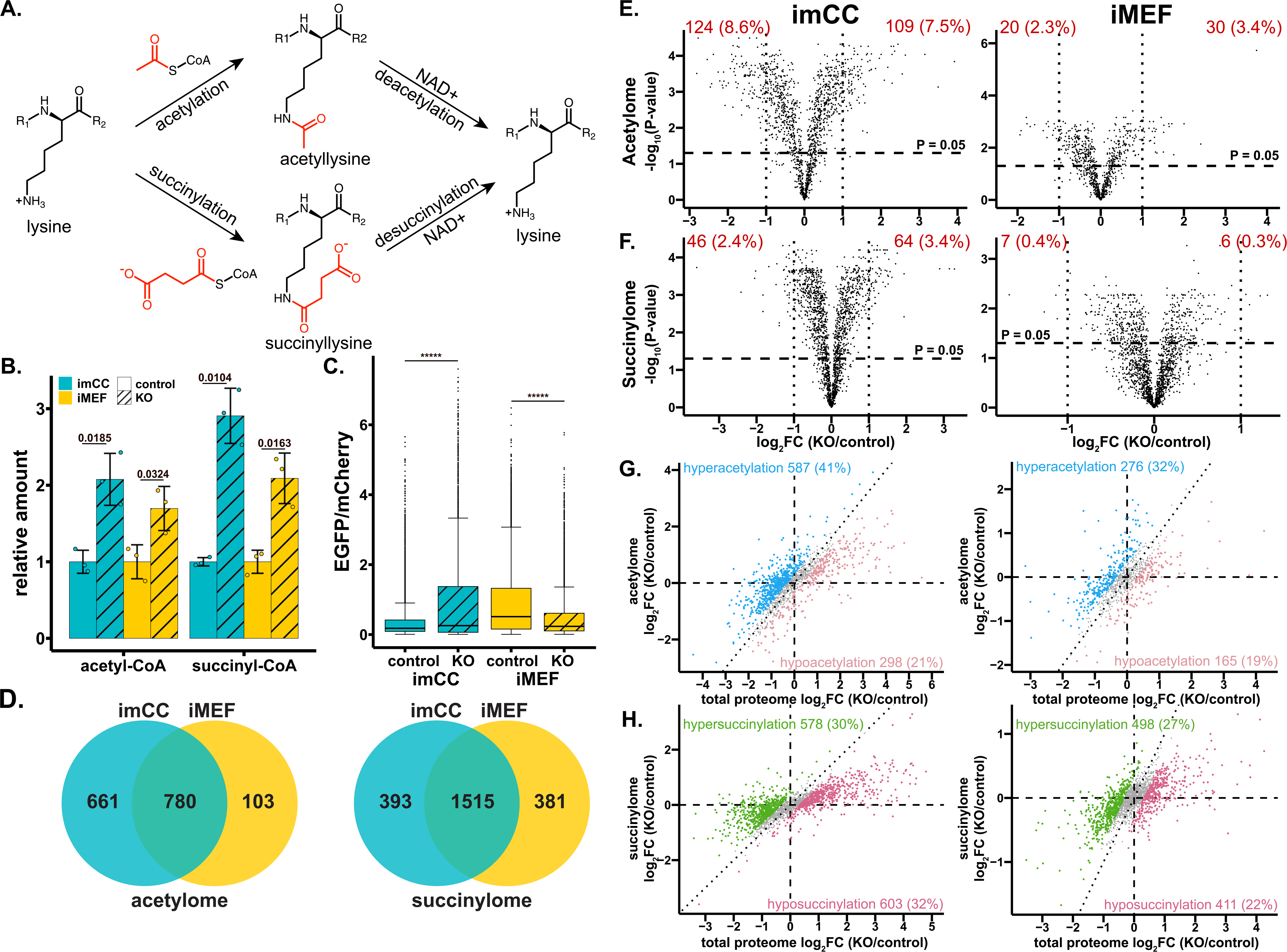
imCCs and iMEFs undergo global disproportionate hyperacetylation but mixed succinylation effects upon SDH loss. **A.** Schematic of lysine acetylation by acetyl-CoA and lysine succinylation by succinyl-CoA, and enzymatic deacylation typically requiring NAD^+^ as a cofactor. **B.** Levels of acetyl-CoA and succinyl-CoA in imCCs and iMEFs measured by LCMS (mean ± SD, *n* = 3 biological replicates per condition). P-values were calculated using two-tailed Welch’s t-tests. **C.** Sirtuin deacetylase activity as measured by activation of acetylation-quenched EGFP relative to mCherry expression in live (DAPI-) cells on flow cytometry. P-values were calculated using Mann-Whitney U-tests. ***** indicates p < 0.00001. **D.** Venn diagram of all identified acetylated and succinylated proteins as shared or unique to imCC and iMEF acylomes. **E-F.** Volcano plots for effects of SDH loss on **E.** acetylome and **F.** succinylome in imCCs and iMEFs. Data are not normalized to total change in protein level. Vertical dotted lines indicate FC of 2. The number of significantly changed (P (FDR-adjusted p-value) < 0.05, |FC| > 2) proteins are labeled in red. **G-H.** Correlation plots between SDH-loss effects on total proteome and **G**. acetylome or **H.** succinylome in imCCs and iMEFs. Dotted diagonal line shows y=x where there is a 1:1 correlation of total proteome and acylome. Proteins with acylation/total protein > 1.25 are colored in blue (disproportionate hyperacetylation) or green (disproportionate hypersuccinylation) and total protein/acylation > 1.25 are colored in light pink (disproportionate hypoacetylation) or dark pink (disproportionate hyposuccinylation) with total counts and percentages per category labeled.

Given the highly reactive nature of acyl-CoA metabolites, we predicted that SDH-loss cells would exhibit global lysine hyperacetylation and hypersuccinylation, particularly in the case of SDH-loss iMEFs that display lower sirtuin activity. We therefore performed immune-enriched TMT-based proteomics to map and quantitate acetyllysine and succinyllysine differences upon SDH loss (Supplemental Fig. S1).

We identified 1444 acetylated and 1908 succinylated proteins in imCCs and 883 acetylated and 1896 succinylated proteins in iMEFs (Fig. 5D). 780 acetylated proteins were shared between imCCs and iMEFs and 1515 succinylated proteins were shared between imCCs and iMEFs. Overall, we were surprised to find that there was not a clear trend toward global hyperacylation in SDH-loss cells. Rather, we detected a mix of acylation changes, suggesting complex regulation of these post-translational modifications, presumably including subcellular compartments with different acyl-CoA concentrations and deacylase enzyme activities. SDH-loss imCCs demonstrated slightly fewer proteins that decreased vs. increased in acetylation upon SDH loss, while iMEFs demonstrated slightly more proteins that increased vs. decreased in acetylation upon SDH loss (Fig. 5E). With respect to quantitative assessment of succinylation, SDH-loss imCCs displayed slightly more proteins that increased in succinylation, while SDH-loss iMEFs displayed similar numbers of proteins with increased and decreased succinylation (Fig. 5F).

Overall, SDH loss had less impact on protein acylation in iMEFs, with only 5.7% of acetylated proteins significantly altered upon SDH loss, compared to 16.1% in imCCs, and 0.7% of succinylated proteins significantly altered by SDH loss in iMEFs compared to 5.8% in imCCs.

We assessed the subcellular distribution of identified acetylated and succinylated proteins in imCCs and iMEFs and found nuclear and cytoplasmic proteins more likely to be acetylated and proteins in secreted and endocytic trafficking compartments less likely to be acetylated (Supplemental Fig. S8A-B). The probability of succinylation was found to be higher for cytoplasmic proteins but not different in the other subcellular compartments (Supplemental Fig. S8A-B), which is surprising since mitochondria are expected to be the main source of succinyl-CoA. Together, these data suggest that more acetylation occurs in the nucleus and cytoplasm and more succinylation occurs in the cytoplasm compared to other cellular compartments.

We wished to perform IPA to analyze the cellular pathways highlighted by proteins whose acylation was significantly altered upon SDH loss. This analysis was possible for imCCs but iMEFs lacked a sufficient census of acylation alterations to enable IPA. Comparison of significantly dysregulated pathways identified by IPA between the total proteome and acetylome of imCCs showed that 17 dysregulated pathways were shared, with 5 upregulated pathways unique to the imCC acetylome (Supplemental Fig. S4D-E), including “citric acid cycle” and “mitotic G1 phase and G1/S transition.” Comparison of dysregulated pathways identified by IPA between total proteome and succinylome in imCCs showed that 17 dysregulated pathways were shared, with 4 dysregulated pathways unique to the imCC succinylome (Supplemental Fig. S4F-G), including extracellular matrix-related pathways. Collectively, these observations suggest hyperacetylation of mitochondrial and nuclear proteins and hypersuccinylation of extracellular matrix proteins upon SDH loss in imCCs, consistent with our original hypothesis.

### SDH loss leads to disproportionate hyperacetylation and mixed disproportionate changes in succinylation

The significant quantitative alterations in total proteomes of imCCs and iMEFs upon SDH loss affect the amount of each protein available for acylation. We therefore focused on acylation changes that were *disproportionate* to changes in the level of each protein upon SDH loss, implying regulation of acylation beyond simple mass action. Both imCCs and iMEFs showed much more disproportionate hyperacetylation upon SDH loss compared to hypoacetylation (Fig. 5G) while both showed similar numbers of disproportionately hypersuccinylated and hyposuccinylated proteins upon SDH loss (Fig. 5H. These data demonstrate that both imCCs and iMEFs display greater disproportionate proteome hyperacetylation upon SDH loss, while alterations in succinylation are relatively balanced.

We performed DAVID functional annotation of disproportionately acylated proteins in SDH-loss imCCs and iMEFs to identify affected pathways and compartments. Disproportionate hyperacetylation in SDH-loss imCCs highlighted nuclear proteins, particularly pathways involving translation and mRNA processing among the top three clusters (Supplemental Table S2). While analysis of disproportionately hyperacetylated proteins in iMEFs upon SDH loss also showed enrichment in translation and mRNA processing among the top three clusters, there was also enrichment of mitochondrial clusters, particularly TCA cycle proteins. In surprising contrast, a mitochondrial cluster was enriched among the top three clusters of disproportionately *hypo*acetylated proteins in imCCs upon SDH loss, as well as an actin cytoskeleton cluster, with two hypoacetylated histones (H4, H2 type 2-A) also driving inclusion of nucleosome and DNA-repair related pathways as affected.

Analysis of disproportionately hypoacetylated proteins in iMEFs upon SDH loss showed enrichment in the actin cytoskeleton and peroxisomal fatty acid metabolism among the top three clusters (Supplemental Table S2). DAVID analysis of disproportionately hypersuccinylated proteins in imCCs and iMEFs upon SDH loss both showed similar enrichment of pathways involving translation and mRNA processing among the top three clusters, while analysis of disproportionately hyposuccinylated proteins showed enrichment of mitochondrial and the actin cytoskeleton clusters among the top three clusters in both imCCs and iMEFs, as well as a cluster with both mitochondrial and peroxisomal fatty acid metabolism only in iMEFs. Thus, while many patterns of altered protein acylation upon SDH loss were similar between imCCs and iMEFs, mitochondrial acetylation patterns were strikingly different between the two cell types.

### SDH loss differentially alters protein acylation across subcellular compartments

Because TCA-cycle-derived acetyl-CoA and succinyl-CoA are produced in mitochondria, we hypothesized that mitochondrial proteins would demonstrate the greatest hyperacylation upon SDH loss. Indeed, mitochondrial proteins experienced hyperacetylation upon SDH loss in both imCCs and iMEFs (Fig. 6A), but, interestingly, mitochondrial hypersuccinylation was detected only in imCCs and not in iMEFs (Fig. 6B). Accounting for total protein levels, only iMEFs showed greater disproportionate hyperacetylation of mitochondrial proteins upon SDH loss (Fig. 6C). Enrichment was observed for acetylated TCA cycle and ETC proteins (Supplemental Fig. S9A,B), consistent with our DAVID functional annotation of disproportionately hyperacetylated proteins in iMEFs. Surprisingly, mitochondrial proteins demonstrated disproportionate *hypo*succinylation in imCCs upon SDH loss (Fig. 6D), including disproportionate hyposuccinylation of TCA cycle and ETC proteins (Supplemental Fig. S9A,B). In contrast, SDH-loss iMEFs exhibited disproportionate hypersuccinylation of ETC proteins (Supplemental Fig. S9B).

**Figure 6.**
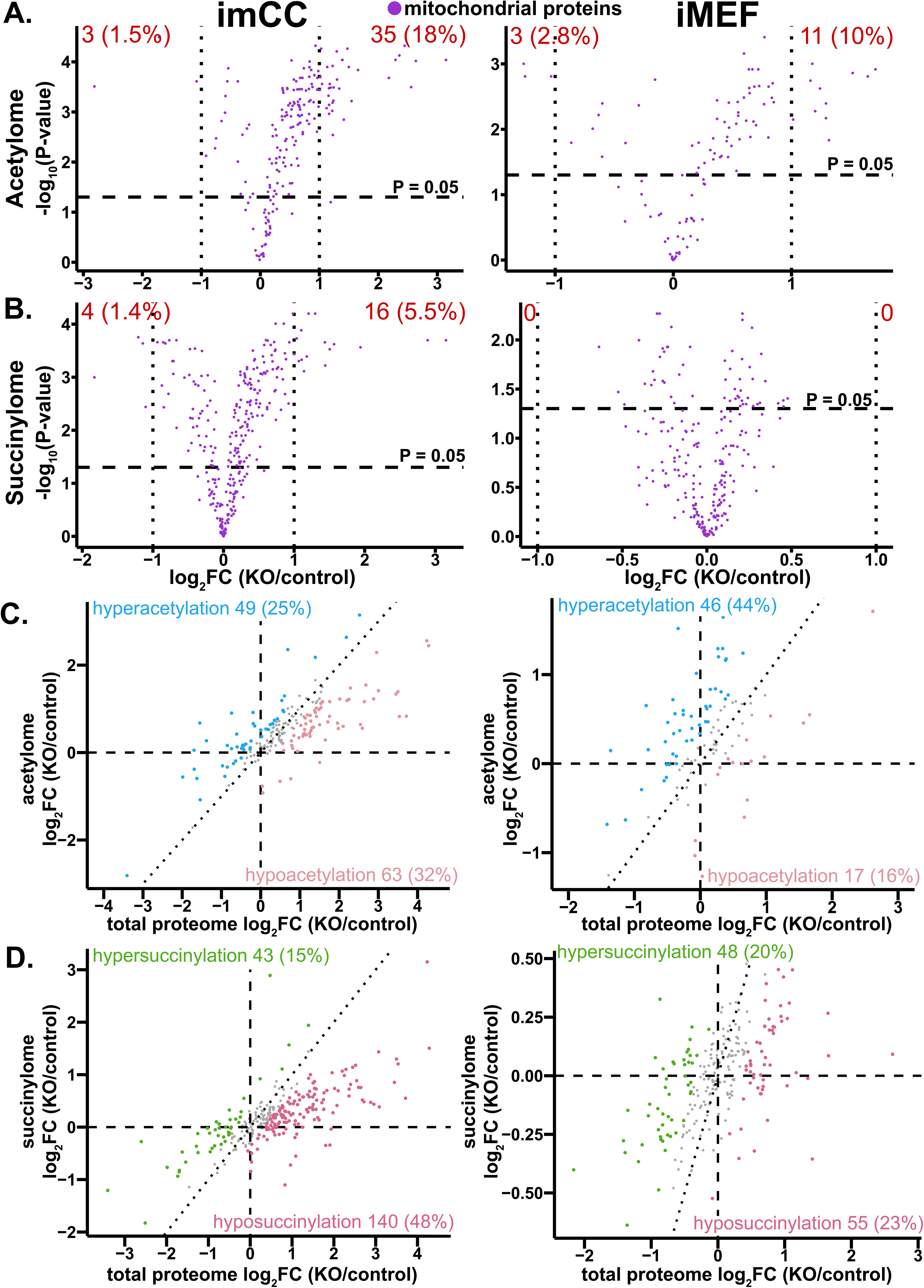
SDH loss induces disproportionate mitochondrial protein hypoacetylation and hyposuccinylation in imCCs but not in iMEFs. **A-B.** Volcano plots for mitochondrial proteins in **A.** acetylome and **B.** succinylome in imCCs and iMEFs. Data are not normalized to total change in protein level. Vertical dotted lines indicate FC of 2. The number and percentage of significantly changed (P (FDR-adjusted p-value) < 0.05, |FC| > 2) proteins are labeled in red. **C-D.** Correlation plots for disproportionate acylation changes in mitochondrial proteins between total proteome and **C.** acetylome or **D.** succinylome in imCCs and iMEFs. Dotted diagonal line shows y=x where there is a 1:1 correlation of total proteome and acylome. Proteins with acylation/total protein > 1.25 are colored in blue (disproportionate hyperacetylation) or green (disproportionate hypersuccinylation) and total protein/acylation > 1.25 are colored in light pink (disproportionate hypoacetylation) or dark pink (disproportionate hyposuccinylation) with total counts and percentages per category labeled.

Because acetyl-CoA and succinyl-CoA acylate histones and other nuclear proteins with epigenetic consequences (22, 47), we assessed if SDH-loss-induced increases in acetyl-CoA and succinyl-CoA were reflected by increased nuclear protein acylation upon SDH loss. Surprisingly, SDH loss was accompanied by hypoacetylation of nuclear proteins in imCCs, but not in iMEFs (Fig. 7A), while similar numbers of proteins showed increased and decreased succinylation in SDH-loss imCCs and iMEFs (Fig. 7B). Accounting for total protein levels, we found disproportionate hyperacetylation of nuclear proteins upon SDH loss in both imCCs and iMEFs, with a four-fold disproportionate hyperacetylation in imCCs compared to two-fold disproportionate hyperacetylation in iMEFs (Fig. 7C). Nuclear proteins also showed disproportionate hypersuccinylation upon SDH loss in both imCCs and iMEFs, with three-fold more hypersuccinylated than hyposuccinylated proteins in imCCs compared to two-fold more in iMEFs (Fig. 7D). Notably, disproportionate histone hyperacetylation upon SDH loss was profound in imCCs but not in iMEFs (Fig. 7E), while more disproportionate histone hyposuccinylation was observed in iMEFs than in imCCs (Fig. 7F). Interestingly, expression of histone acetyltransferases was significantly decreased upon SDH loss in imCCs but not iMEFs (Supplemental Table S3), perhaps an adaptive response to histone hyperacetylation.

**Figure 7.**
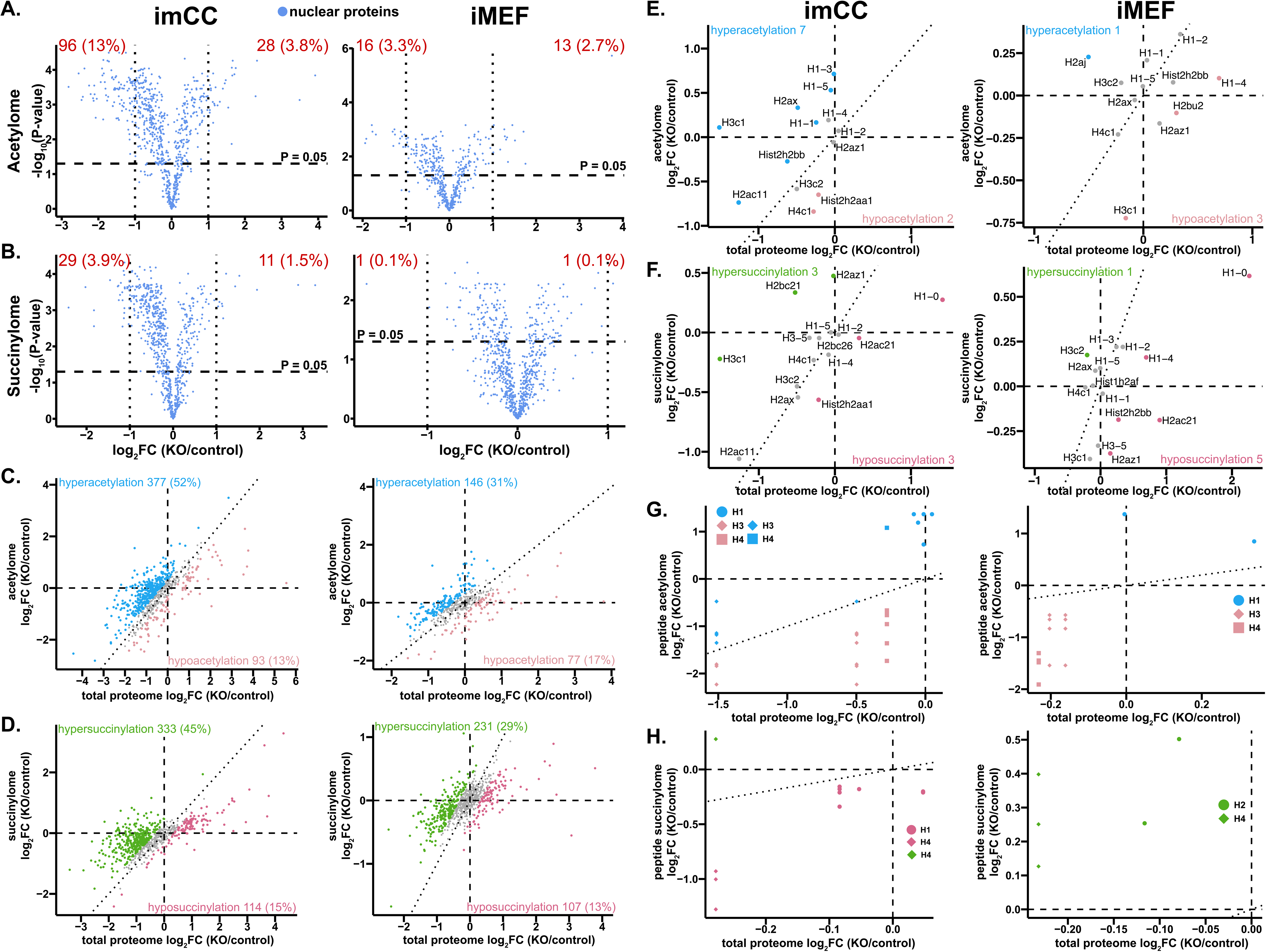
SDH loss leads to disproportionate nuclear protein hyperacetylation and hypersuccinylation in both imCCs and iMEFs. **A-B.** Volcano plots for nuclear proteins in **A.** acetylome and **B.** succinylome in imCCs and iMEFs. Data are not normalized to total change in protein level. Vertical dotted lines indicate FC of 2. The number and percentages of significantly changed (P (FDR-adjusted p-value) < 0.05, |FC| > 2) proteins are labeled in red. **C-D.** Correlation plots for nuclear proteins between total proteome and **C.** acetylome or **D.** succinylome in imCCs and in iMEFs. **E-F.** Correlation plots for the indicated histones between total proteome and **E.** acetylome or **F.** succinylome in imCCs and iMEFs. **G-H.** Correlation plots for the indicated histones peptides between total proteome and **E.** peptide acetylome or **F.** peptide succinylome in imCCs and iMEFs. Dotted diagonal line shows y=x where there is a 1:1 correlation of total proteome and acylome. Proteins with acylation/total protein > 1.25 are colored in blue (disproportionate hyperacetylation) or green (disproportionate hypersuccinylation) and total protein/acylation > 1.25 are colored in light pink (disproportionate hypoacetylation) or dark pink (disproportionate hyposuccinylation) with total counts and percentages per category labeled.

imCCs and iMEFs showed different patterns of histone H2A and H2B expression and acylation (Supplemental Table S4). Both expression and acylation of H2A types 1-F and 2-C were only detected in iMEFs, while acylation of H2A type 2-A and H2B type 2-E was only detected in imCCs. Both expression and acylation of H2A type 1-G, H2A.V, H2B type 3-B were only detected in imCCs.

To assess in greater detail the impact of SDH loss on histone acylation, we mapped changes in acetylated and succinylated histone peptides. We identified a total of 236 histone acetylation and succinylation sites, of which 67 were solely acetylation sites, 108 were solely succinylation sites, and 61 sites were detected in both acetylated and succinylated forms (Supplemental Table S5). Intriguingly, acetylation and succinylation sites tended to be located in different histone domains. Acetylation sites predominantly mapped to histone tails (63%) while succinylation sites predominantly mapped to histone bodies (70%). This pattern was observed for both cell types. Because histone acylation modifications can impact chromatin regulation through both direct electrostatic effects and by recruitment of epigenetic readers, these data are consistent with histone tail acetylation affecting epigenetic control more through the latter mechanism, while effects of histone body succinylation may involve the former mechanism (48).

We then focused on histone peptides in both cell types that displayed significantly altered acylation (p<0.05) upon SDH loss. H1 peptides were disproportionately hyperacetylated in both imCCs and iMEFs upon SDH loss, while H3 and H4 peptides tended to be disproportionately hypoacetylated upon SDH loss (Fig. 7G). Conversely, H1 and H4 peptides were disproportionately hyposuccinylated in SDH-loss imCCs, while H3 and H4 peptides tended to be disproportionately hypersuccinylated upon SDH loss in iMEFs (Fig. 7H), in agreement with our previous studies (22). Together, these results suggest that, on a peptide level, disproportionate histone acetylation changes upon SDH loss are relatively similar between imCCs and iMEFs while disproportionate succinylation changes differ.

In contrast to the observed skewed acylation changes in mitochondria and nucleus upon SDH loss, cytoplasmic proteins showed a more balanced census of increased and decreased acetylated and succinylated proteins upon SDH loss in both imCC and iMEFs (Fig. 8A, B). After accounting for total protein levels, both imCCs and iMEFs demonstrate disproportionate cytoplasmic protein hyperacetylation and hypersuccinylation upon SDH loss, consistent with our original hypothesis (Fig. 8C, D).

**Figure 8.**
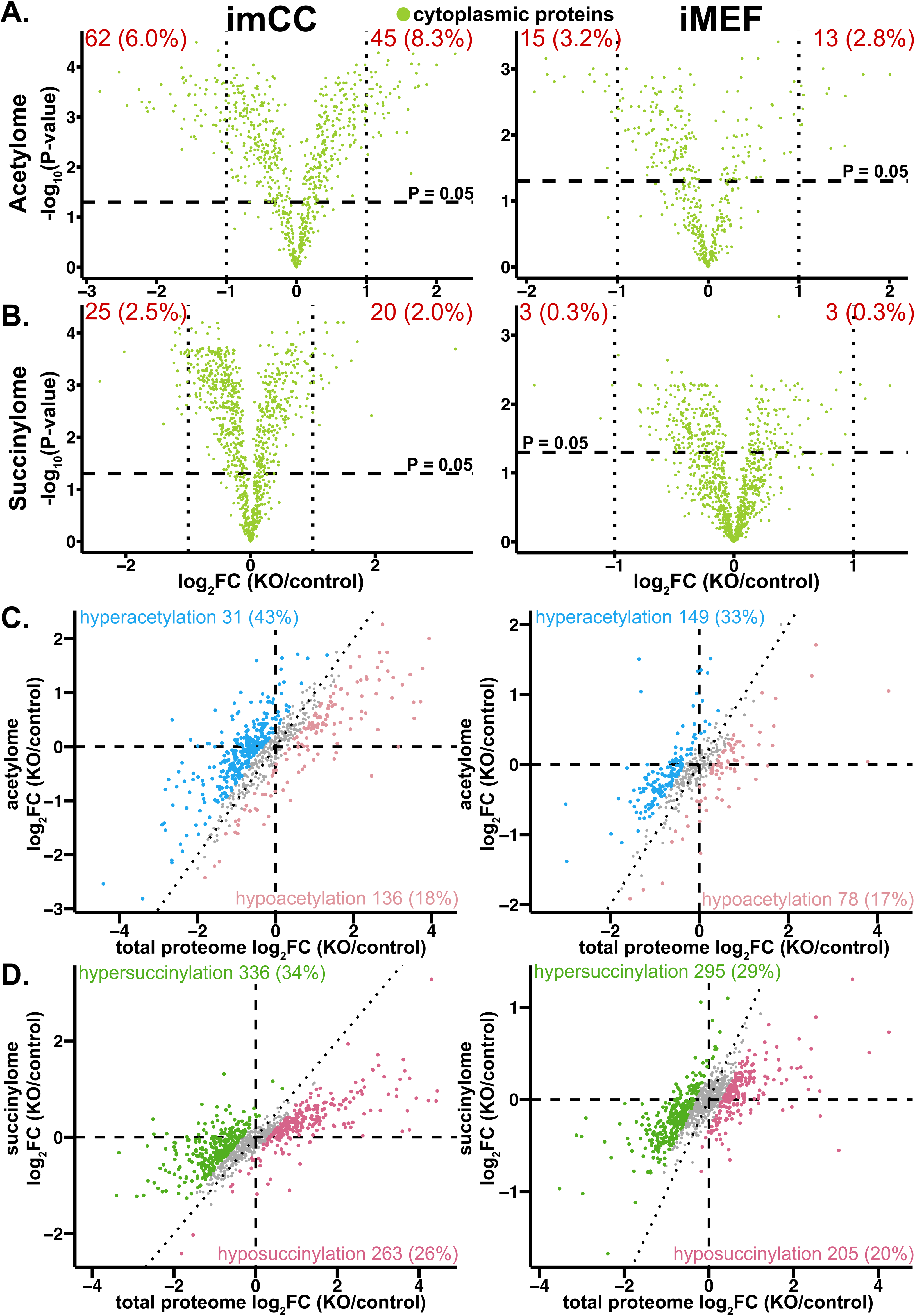
SDH loss leads to disproportionate cytoplasmic protein hyperacetylation and hypersuccinylation in both imCCs and iMEFs. **A-B.** Volcano plots for cytoplasmic proteins in **A.** acetylome and **B.** succinylome in imCCs and iMEFs. Data are not normalized to total change in protein level. Vertical dotted lines indicate FC of 2. The number and percentages of significantly changed (P (FDR-adjusted p-value) < 0.05, |FC| > 2) proteins are labeled in red. **C-D.** Correlation plots for cytoplasmic proteins between total proteome and **C.** acetylome or **D.** succinylome in imCCs and in iMEFs. Dotted diagonal line shows y=x where there is a 1:1 correlation of total proteome and acylome. Proteins with acylation/total protein > 1.25 are colored in blue (disproportionate hyperacetylation) or green (disproportionate hypersuccinylation) and total protein/acylation > 1.25 are colored in light pink (disproportionate hypoacetylation) or dark pink (disproportionate hyposuccinylation) with total counts and percentages per category labeled.

## Discussion

The etiology of PPGL tumors remains mysterious on multiple levels. For example, while disruptions to the TCA cycle are likely to be tumorigenic due to oncometabolite accumulation, the basis for the cell-type specificity of induced tumorigenesis is unclear. Although both SDH-loss and VHL-loss tumors share chronic activation of HIF transcription factors and pseudohypoxia, SDH-loss tumors are more aggressive tumors than those with VHL loss. Here, we demonstrate through detailed quantitative analysis that SDH loss significantly alters the proteomes and acylproteomes of two available immortalized mouse cell line models of SDH-loss tumors. Profound energy deficiency occurs in both imCCs and iMEFs upon SDH loss, yet only SDH-loss imCCs show upregulation of mitochondrial TCA cycle and FAO proteins with increased FAO activity. Despite accumulation of lysine-reactive acetyl-CoA and succinyl-CoA in both SDH-loss cell models, proteins are generally disproportionately hyperacetylated but, surprisingly, show mixed changes in succinylation levels upon SDH loss.

### An imperfect comparison

Our studies were designed to examine how cell context influences the response to SDH loss in the context of PPGL tumorigenesis. The field offers a paucity of cell and animal models to address this question. The two paired mouse cell models we selected for study are derived from the adrenal medulla (imCCs) and embryonic fibroblasts (iMEFs) and have been previously studied as models of SDH-loss PPGL. For historical reasons, the genetic defects in the two models affect different subunits of the SDH complex: SDHB is deleted in imCCs while SDHC is deleted in iMEFs. Since deficiency in any SDH subunit or its assembly factors results in loss of SDHB (Supplemental. Fig. S7B) with corresponding loss of SDH enzymatic function (45, 49–54), loss of either SDHB or SDHC should be functionally equivalent at the enzymatic level in cell culture. However, we recognize that idiosyncratic variations in the properties of the remaining undegraded SDH sub-complexes could contribute to additional differences between SDHB-loss imCCs and SDHC-loss iMEFs, potentially confounding simple attribution of observed differences to cell type. This caveat must be born in mind. Resolving these differences will require future development of truly matched control and SDH-loss cell models affected by identical SDH subunit defects across multiple cell types. Such an analysis is beyond current available models.

### Fundamental low energy metabolism in SDH-loss cells

The concept of metabolic defects driving tumorigenesis challenges the classic cancer paradigm of rapid, abnormal cellular proliferation driven by oncogenes or loss of tumor suppressor genes regulating the cell cycle. In particular, perturbation of the TCA cycle and ETC are expected to negatively impact energy homeostasis. The markedly slower growth of SDH-loss cells and other features (e.g., resistance to lentiviral infection) have long suggested to us that these cells suffer from a strikingly impaired metabolism with accompanying energy stress. Indeed, we now report for the first time that both SDH-loss imCCs and iMEFs show profound energy deficits with both low ATP/ADP and high NAD^+^/NADH ratios (Fig. 4C,E). The high NAD^+^/NADH ratio reflects a more oxidizing state and indicates a deficiency in ETC electron donors for ATP production, likely contributing to the observed low ATP/ADP ratio and restricting SDH-loss cell proliferation even in rich media. Elevated NAD^+^/NADH ratios have also been reported for other cell models of SDH loss (HEK293T, 143B) when Complex I function is retained but not when Complex I function is lost (55). This pattern is consistent with the presence of residual Complex I activity in SDH-loss imCCs (36, 37), despite significant morphological pathology detected in ultrastructural features of mitochondria in both SDH-loss imCCs and iMEFs (37). However, SDH-loss iMEFs that lack Complex I activity must have a different mechanism of NAD^+^ regeneration. While the low ATP/ADP energetic stress in SDH-loss cells presumably drives a compensatory shift towards glycolysis (i.e., the Warburg effect), we showed previously that only SDH-loss iMEFs, but not imCCs, displayed increased glycolysis with pyruvate fermentation to lactate (37). Together, these results suggest that SDH-loss imCCs display high NAD^+/^NADH due to residual Complex I activity in the ETC, whereas SDH-loss iMEFs display high NAD^+/^NADH due to pyruvate fermentation to lactate. Neither approach is sufficient to restore normal ATP/ADP ratios, creating energy stress and limiting the rate of SDH-loss cell growth.

It is noteworthy that solid tumors also have been observed to downregulate energy-intensive processes such as tissue-specific protein synthesis in the context of low ATP production and reduced TCA cycle flux in the case of mouse pancreatic cancer models (56). Similarly, we find that both imCCs and iMEFs significantly downregulate translation machinery and related protein processing upon SDH loss. Impaired ribosomal biogenesis is known to impact the cell cycle through downregulation of Cdk6 (57, 58), which we find to be decreased by five-fold in both imCCs and iMEFs upon SDH loss. These findings may explain the accumulation of cells in G1 phase and a decreased S phase population in the cell cycles of SDH-loss imCCs and iMEFs (37). Ribosomal deficiency can also lead to higher levels of p53 (58), which we find to be increased 6-fold in SDH-loss iMEFs but decreased 10-fold in SDH-loss imCCs. Because similar cell cycle changes are observed upon SDH loss in both imCCs and iMEFs (37), the implications of this striking difference in p53 levels between the SDH-loss cell lines is unclear.

Together, our findings indicate that energy deficiency and downregulation of energy-intensive processes such as ribosomal biogenesis are shared features of SDH-loss cells, though compensatory mechanisms resulting in an increased NAD^+^/NADH ratio differ between imCCs and iMEFs. These observations are consistent with the substantially reduced growth rate of SDH-loss cells, a paradoxical behavior if SDH-loss imCCs are considered models of SDH-loss PPGL tumors. The emerging picture of SDH loss is one of fundamentally disabled cells struggling to grow under severe energy deficiency, yet still able to survive, with possible resistance to apoptosis (59), and eventually proliferate, albeit slowly, into tumors.

### Increased FAO in SDH-loss imCCs

While glycolysis and the TCA cycle are typically the main sources of electrons carried by NADH to Complex I of the ETC, other catabolic processes such as FAO and amino acid catabolism can generate high-energy electron equivalents as NADH or FADH_2_, especially in the setting of TCA cycle and ETC disruption (60). We now report that SDH-loss imCCs, but not iMEFs, accumulate FAD and upregulate FAD-dependent enzymes, such as acyl-CoA dehydrogenases involved in FAO and the electron transfer flavoprotein (ETF) complex that funnels electrons from at least 14 different flavoenzymes into the ETC (61). SDH-loss imCCs are also characterized by a 2.5-fold increase in cytoplasmic FAD synthase, likely a compensatory response to increased FAD demand from flavoenzymes in FAO, BCAA catabolism, and choline catabolism, all of which are upregulated in imCCs but not iMEFs upon SDH loss. Increased FAD synthase expression has also been observed in pancreatic cancer models (62), suggesting that upregulation of FAD-dependent catabolic pathways may be an important alternative source of energy for abnormally proliferating cells that retain ETC functionality. Upregulation of BCAA and choline catabolism, along with fatty acid metabolism, TCA cycle, oxidative phosphorylation, and extracellular matrix components have also been found in *SDHB*-mutant head and neck paragangliomas compared to normal nerve tissue (63). We find all these characteristics in cultured SDH-loss imCCs, suggesting that SDH-loss imCCs indeed better recapitulate aspects of human SDH-loss PPGL tumors than do SDH-loss iMEFs. While SDH-loss imCCs show only mild dependence on FAO under our nutrient-rich culture conditions (Fig. 4K), further study could determine the importance of FAO for SDH-loss cell growth under more physiologically relevant conditions.

### Complex pattern of altered protein acylation upon SDH loss

We originally hypothesized that SDH-loss cells would be characterized by profound global protein hyperacylation due to accumulation of lysine-reactive acyl-CoAs upstream of SDH in the TCA cycle. Our results indicate a much more nuanced picture, wherein alterations in protein acylation upon SDH loss are complex despite elevated levels of acetyl-CoA and succinyl-CoA. Changes in sirtuin deacylase activity (perhaps reflecting increased levels of sirtuin cofactor NAD^+^) undoubtedly play roles in this complexity. In fact, both disproportionate protein hyperacylation and hypoacylation are observed, depending on the protein (and even the protein domain), the cell type, and the subcellular compartment. Accumulation of succinyl-CoA upon SDH loss is consistent with previous studies (22) and is expected to occur as the metabolite precursor to succinate in the TCA cycle. While intermediates upstream of succinyl-CoA do not necessarily accumulate in SDH loss (37), accumulation of acetyl-CoA may result from increased FAO and increased glycolytic flux, which has been found to be required for maintenance of H3K27ac in breast cancer (64). Altered localization and flux of acetyl-CoA between cellular compartments has also been found to impact protein acetylation and regulate FAO and mitochondrial metabolism (65). Additionally, significant upregulation of the cholesterol synthesis/mevalonate pathway in iMEFs but not imCCs upon SDH loss (Fig. 1E,F) may provide an alternative means for cytosolic acetyl-CoA consumption, decreasing the fraction of cytoplasmic proteins undergoing disproportionate hyperacetylation in iMEFs compared to imCCs upon SDH loss (Fig. 8C). Changes in levels of site-specific protein acylation affecting non-histone proteins have previously been shown to impact protein function and stability (23, 27, 29, 66) and may impact tumor growth and immune responses (25, 26, 31, 67). While assessment of the growth signaling impacts of individual acylation sites is beyond the scope of this work, SDH loss-induced alterations in protein acylation could have tumorigenic consequences that warrant further study.

### Altered histone acylation and potential epigenetic effects

Histone acetylation is a well-studied modification that impacts transcription both by directly weaking nucleosomes through reduced electrostatic interactions between histones and with DNA, and through recruitment of epigenetic readers (47). More recently, histone succinylation has also been shown to impact transcription, likely through nucleosome destabilization by electrostatic repulsion of negatively charged succinyllysines (22, 48, 68). Our findings that acetylation sites are enriched in histone tails relative to histone bodies while succinylation sites are enriched in histone bodies relative to histone tails support a model in which acetylation marks act more through reader recruitment while succinylation marks act more through fundamental electrostatic effects (Supplemental Table S4). While on a protein level SDH-loss imCCs demonstrate disproportionate histone hyperacetylation and disproportionate histone hyposuccinylation (Fig.7E, F), disproportionate acylation also varied by peptide (Fig. 7G-H), suggesting that regulation of site-specific histone acylation is highly nuanced. Additional studies are needed to determine the functional consequences of alterations in site-specific histone acylation and patterns of histone acylation in the setting of SDH loss.

## Conclusions

Together, the results of this work demonstrate that SDH loss imposes a profound energy-deficient state, with significant, cell type-specific changes affecting the proteomes and acylproteomes in two cell models (Fig. 9). This work reveals cell type-specific adaptations to the energy deficit experienced by SDH-loss cells and suggests that SDH-loss imCCs possess superior metabolic plasticity, thereby providing clues to how SDH loss may be selectively tolerated in certain tissues.

**Figure 9.**
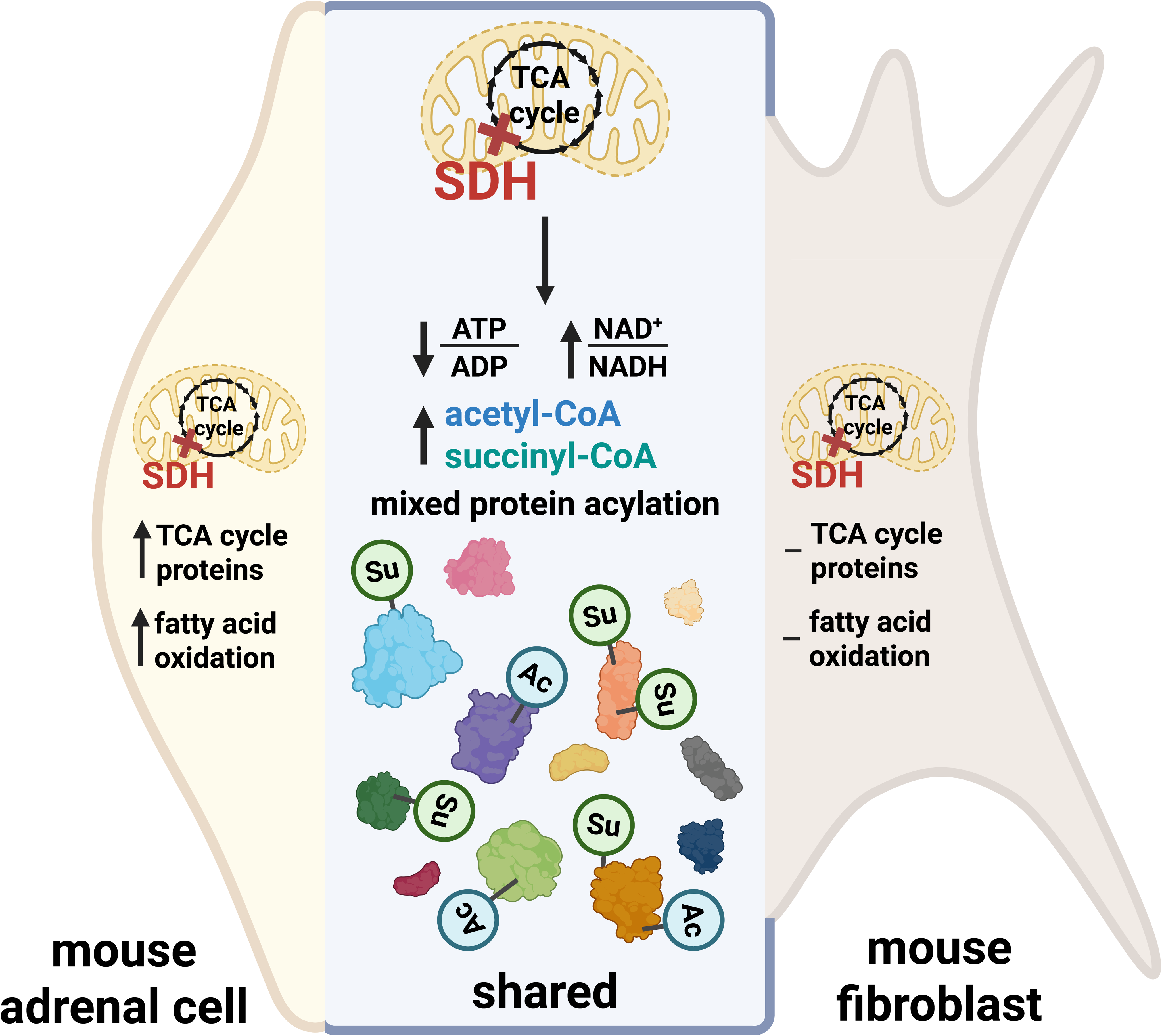
Graphical abstract depicting the metabolic and proteomic impact of SDH loss on imCCs and iMEFs.

## Materials and methods

### Cell culture

imCCs (35) and iMEFs (34) were maintained in high glucose Dulbecco’s modified Eagle’s medium (DMEM) (Gibco 11965) supplemented with 10% heat-inactivated fetal bovine serum (FBS) (Gibco A5669801), 5% penicillin-streptomycin (Gibco 15140122), 1 mM GlutaMAX™ (Gibco 35050061), 1 mM sodium pyruvate (Gibco 11360070), 10 mM HEPES buffer (Gibco 15630080) and 1× MEM non-essential amino acids [100 µM final concentration each of glycine, alanine, asparagine, aspartic acid, glutamic acid, proline, and serine (Gibco 11140050)], referred to as standard culturing media. For assessment of proliferation in modified media, the following nutrient adjustments were made to the standard culturing media, based on DMEM without glucose, glutamine, or phenol red (Gibco A1443001): reduced glucose (5 mM from 25 mM) and reduced GlutaMAX (0.55 mM from 5 mM) approximating physiological amounts found in human plasma. Cells were adapted to the modified media for at least two doublings before being used for subsequent assays. All cells were cultured at 37°C, 95% humidity in room air (21% O_2_) with 5% CO_2_.

### Proliferation rate determination

Cells were seeded at 10,000 cells per well in 12-well plates and cultured overnight. Triplicate wells were then counted to obtain starting cell counts and remaining triplicate wells received fresh media containing the indicated treatments. Final cell counts were obtained after three doublings for each cell type and proliferation rates were calculated using equation 1 below. Cells were counted using a Luna II cell counter (Logos Biosystems).

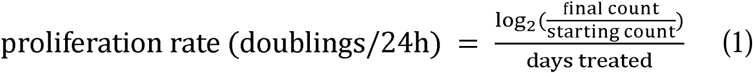

### Sirtuin activity assay

Sirtuin activity was measured using a dual-fluorescence reporter as previously described (34). Cells were seeded in 6-well plates and cultured overnight. Cells were then co-transfected with pcDNA-mCherry-EGFP(K85AcK) and pCMV-AcKRS-tRNAPyl (gifts of Peter Schultz, Scripps Research Institute) using FuGENE HD (Promega) and Opti-MEM (Gibco). 5 mM acetyllysine (AcK) was added 1 h after transfection. EFP and mCherry fluorescence intensity was quantified by flow cytometry 30 h after transfection. Live (DAPI-) single cells were gated for mCherry and EGFP(K85AcK) expression analysis. Relative sirtuin activity was quantified as the degree of EGFP fluorescence relative to mCherry fluorescence in dual EGFP-mCherry positive cells after subtraction of background signal in each channel for each cell type.

### Sample processing and TMT peptide labeling

SDH-loss and control imCCs and iMEFs were washed with cold PBS before scraping into 1.5 mL lysis buffer (20 mM triethyl ammonium bicarbonate (TEAB), pH 8.5, 8 M urea, 1× protease inhibitors (Pierce A32963)) per sample and frozen at -80°C. Samples were thawed and probe sonicated for lysis and protein extraction. Protein concentration was determined by BCA Assay (Pierce 23225) and aliquots of 2 mg protein were reduced at room temperature for 1 h with 10 mM of tris-(2-carboxyethyl)phosphine followed by alkylation for 20 min in the dark at room temperature with 20 mM iodoacetamide. A sufficient volume of 20 mM TEAB, pH 8.5 buffer were added to reduce the urea concentration to 0.9 M and 100 µg of trypsin (1:20 enzyme to substrate ratio, w/w, Worthington) was added to the samples before incubating at 37°C overnight. The digestion reaction was quenched using 20% trifluoroacetic acid (TFA) and desalted with using Sep-Pak C_18_ light cartridges (Waters) using 0.1% TFA / acetonitrile eluting solvent. Eluted peptides were lyophilized, solubilized in 300 mM HEPES, pH 8.5 buffer and labeled with 2 mg TMT sixplex or 10plex reagents (Thermo Fisher Scientific 90061, 90110) in acetonitrile for 1 h at room temperature and quenched with 5% hydroxylamine. 5 µL aliquots from each labeling reaction were combined and analyzed by nanoLC-tandem mass spectrometry to confirm >99% labeling efficiency. The remainder of the reactions were pooled by normalizing with reporter ion intensities to combine equal amounts of each sample. Excess TMT reagent was removed from the mixture using a Sep-Pak C_18_ Plus cartridge (Waters) with 0.1% TFA / acetonitrile eluting solvent and the eluted peptides were lyophilized.

### Immunoaffinity enrichment of lysine acetyllysine and succinyllysine peptides

TMT-labeled acetylated and succinylated peptides were purified using PTMScan HS kits (Cell Signaling Technologies 46784, 60724) following the manufacturer’s recommended protocol. Briefly, for each enrichment, approximately 2 mg of lyophilized TMT-labeled peptides were solubilized in HS IAP binding buffer and incubated with 20 µL of anti-acetyllysine or anti-succinyllysine magnetic bead slurry for 2 h at 4°C. After wash steps, peptides were eluted from the beads with IAP elution buffer into autosampler vials and dried on a SpeedVac concentrator.

### Tandem mass spectrometry acquisition of TMT-labeled acetylated and succinylated peptides

Acetylated and succinylated TMT-labeled peptides were analyzed using an Orbitrap Eclipse mass spectrometer (Thermo Fisher Scientific) coupled to a Vanquish Neo UHPLC system (Thermo Fisher Scientific) with 0.1% formic acid in 98% water / 2% acetonitrile for solvent A and 0.1% formic acid in 80% acetonitrile / 10% isopropanol / 10% water for solvent B. The dried peptides were solubilized in 0.2% TFA / 0.0005% Zwittergent 3-16 and pumped onto a Halo C18 2.7 µm EXP stem trap (Optimize Technologies, Oregon City, OR) with solvent A at a flow rate of 8 µL/min. The trap was placed in line with a PepSep C18 2.7 µm, 40 cm × 100 µm column (Bruker) and the peptides were separated at a flow rate of 350 nL/min with a gradient of 4% B to 45% B over 154 min, then 45% B to 90% B over 12 min and hold at 90% B for 5 min. Data were acquired by data-dependent acquisition with a 3 s cycle time. The MS1 survey scan range was from 360-1600 m/z at resolution 120,000 (at 200 m/z) and the AGC set for a maximum of 1E6 ions and a 50 ms ion injection time. Ions in the scan range of 360-1600 m/z with positive charge states from 2-5 were sequentially selected with an isolation width of 0.7 m/z for high energy collisional dissociation (HCD) fragmentation at a NCE setting of 38. The MS2 Orbitrap scans were at resolution 50,000 with the AGC setting at 250% (2E5 ions) and the maximum ion injection time was set to 200 ms. The dynamic exclusion feature was used to prevent ions selected for MS2 and any ions within an m/z of 7 ppm from being selected for fragmentation for 30 s.

### Modified TMT-labeled peptide identification and quantification

Mass spectrometry raw data files were analyzed using Proteome Discoverer 3.0 (Thermo Fisher Scientific), set for MS2 reporter ion quantification with TMT 10plex isobaric labels. Raw files were searched against a Swiss-Prot mouse (2024_02) database using Sequest HT with parameters set for full trypsin specificity, allowing 3 missed cleavages and the following variable modifications: oxidized methionine and N-terminal protein acetylation, TMT-labeled lysine, and either succinyllysine or acetyllysine. Fixed modifications were carbamidomethyl-cysteine and peptide N-terminal TMT. Mass tolerances were set at 10 ppm for precursor ions and 0.02 Da for MS2 fragment ions. Peptide identifications were filtered at 1% FDR using the Percolator node. Reporter ion channel correction factors were applied to PSMs and filtered to exclude peptides exceeding the threshold maximum of 50% isolation interference. The IMP-ptmRS node was used to assign PTM site probability with the reporting threshold set to 75%. Acetylated and succinylated TMT-labeled peptide and protein results and their matched raw reporter ion intensity values were exported to Excel and group comparisons were made with an in-house R-script. Sample intensity values were normalized by median subtraction and log_2_ transformed and significant differences determined with unpaired, two-tailed t-tests with Benjamini-Hochberg correction. Any proteins with q > 0.01, missing values in any TMT channel, or no acylation sites identified were excluded from downstream analyses.

### Peptide fractionation with basic reverse phase HPLC for total protein

1 mg dried TMT-labeled peptide sample was solubilized in 5% DMSO / 5 mM ammonium formate, pH 8.5 and fractionated using basic pH reverse phase HPLC on an UltiMate 3000 RSLC HPLC system (Thermo Fisher Scientific) with a XBridge Peptide BEH C18, 3.5 µm, 4.6 mm × 250 mm column (Waters 186003570). The mobile phases were 5 mM ammonium formate, pH 8.5 for solvent A and 5 mM ammonium formate, pH 8.5 / 90% acetonitrile/ 10% water for solvent B. A flow rate of 0.5 mL/min was used and gradient conditions were 5% B to 60% B over 60 min, then to 80% B in 2 min and hold for 5 min. A total of 96 fractions were collected over the 80-min period and were concatenated to 24 fractions for nanoLC-tandem orbitrap mass spectrometry analysis.

### NanoLC-tandem mass spectrometry data acquisition for total protein

Peptide fractions were analyzed by nanoLC-tandem mass spectrometry using an Orbitrap Eclipse mass spectrometer (Thermo Fisher Scientific) coupled to a Vanquish Neo UHPLC system (Thermo Fisher Scientific) with 0.1% formic acid in 98% water / 2% acetonitrile for solvent A and 0.1% formic acid in 80% acetonitrile / 10% isopropanol / 10% water for solvent B. Each fraction was solubilized in 0.1% formic acid and pumped onto a Halo C18 2.7 µm EXP stem trap (Optimize Technologies, Oregon City, OR) with solvent A at a flow rate of 8 µL/min. The trap was placed in line with a PepSep C18 2.7 µm, 40 cm × 100 µm column (Bruker) and the peptides were separated at a flow rate of 350 nL/min with a gradient of 4% B to 40% B over 120 min, then 40% B to 90% B over 12 min and hold at 90% B for 5 min. Data were acquired using a real time search MS3 method for TMT quantitation. The MS1 scans were performed in the Orbitrap, with a mass range of 350-1500 m/z, AGC at 100%, 25 ms max IT and resolution at 120,000 at 200 m/z. Ions were selected with an isolation width of 0.7 and fragmented by CID in the ion trap using the turbo scan rate and the MS2 spectra were searched in real time against a Swiss-Prot mouse database with parameters set to allow one missed cleavage, static modifications of carbamidomethyl-cysteine, TMT-labeled lysine and peptide N-terminus, and oxidized methionine as a variable modification. Peptide spectral matches meeting minimum values of 1.2 for Xcorr, 0.1 for dCN and delta mass of 12 ppm or less triggered a MS3 scan event for TMT reporter ion quantitation. Up to 10 fragment ions in the mass range 200-1500 from the matched spectra were selected by synchronous precursor selection to generate the reporter ion MS3 spectra with HCD fragmentation at 55% NCE and orbitrap scanning from 100-500 m/z at 50k resolution. The MS3 AGC setting was 500% with the max ion injection time of 86 ms. Dynamic exclusion was used to prevent ions selected for MS2 and any ions within an m/z of 7 ppm from being selected for fragmentation for 25 s and the close out feature was implemented to restrict the MS3 trigger to a maximum of 6 peptide matches to a protein in each fraction run.

### Total protein identification and quantification

Mass spectrometry raw data files were analyzed using Proteome Discoverer 3.0 (Thermo Fisher Scientific), set for MS3 reporter ion quantification with TMT 10plex isobaric labels. Raw files were searched against a Swiss-Prot mouse (2024_02) database using Sequest HT with parameters set for full trypsin specificity, allowing 2 missed cleavages and the following variable modifications: oxidized methionine and N-terminal protein acetylation. Fixed modifications were carbamidomethyl-cysteine, TMT-labeled lysine and peptide N-terminal TMT. Mass tolerances were set at 10 ppm for precursor ions and 0.5 Da for MS2 fragment ions. Protein identifications were filtered at 1% FDR using the Percolator node. The MS3 TMT reporter ion channel intensities were reported with correction factors applied to PSMs and no filtering with isolation interference set at 100%. The matched proteins and raw reporter ion intensity values were exported to Excel and group comparisons were made with an in-house R-script. The sample intensity values were normalized by median subtraction and log_2_ transformed and significant differences determined with unpaired, two-tailed t-tests with Benjamini-Hochberg correction. Any proteins with q > 0.01 or missing values in any TMT channel were excluded from downstream analyses.

### Secondary proteomics analysis

Principal component analysis (PCA) was performed with prcomp in R.

### DAVID functional annotation

Analysis of functional annotation enrichment for differentially succinylated peptides was performed by extracting the gene identifiers for peptides having succinylation log_2_(fold-change) differing by more than 0.2 from the corresponding peptide abundance log_2_(fold-change). Functional enrichment analysis of functional annotations (UP_KW_BIOLOGICAL_PROCESS, UP_KW_CELLULAR_COMPONENT, UP_KW_MOLECULAR_FUNCTION, UP_KW_PTM), gene ontologies (GOTERM_BP_DIRECT, GOTERM_CC_DIRECT, GOTERM_MF_DIRECT) and pathways (KEGG_PATHWAY, REACTOME) was performed using the DAVID functional annotation tool with default settings and medium stringency (40, 41). Terms were considered significantly enriched if the Benjamini-Hochberg (B-H) corrected p-value < 0.05.

### Ingenuity Pathway Analysis

Significantly changed (FDR adjusted p-value < 0.05 and |fold change| ≥ 2) proteins, acetylated proteins, and succinylated proteins were used as inputs for QIAGEN Ingenuity Pathway Analysis (IPA, (38)). IPA performs enrichment analysis of known canonical pathways in the QIAGEN IPA library using the list of significantly changed proteins and their directionality of change. Canonical pathways were considered statistically significantly altered if the number of significantly changed proteins for a given pathway exceeded the number predicted by the significance criterion. For this study, a threshold Benjamini-Hochberg (B-H) corrected p-value of < 0.05 (-log(B-H p-value) > 1.3) was used. Pathways with a z-score ≥ 2 are predicted to be upregulated and those with a z-score ≤ -2 are predicted to be downregulated.

### Subcellular compartment identification

Subcellular compartment protein lists were downloaded and curated from UniProtKB (01/02/2025). Only reviewed (Swiss-Prot) *Mus musculus* proteins were included in each list and duplicate proteins were removed. Each list was defined as follows: mitochondrion: SL-0173 (1273 proteins); nucleus: SL-0191 (4884 proteins); endoplasmic reticulum and Golgi apparatus (1997 proteins): SL-0095:0099, SL-0132, SL-0235:0237, SL-0239, SL-0248, SL-0266, SL-0326; endosomes/autosomes/lysosomes/phagosomes (1998 proteins): SL-0023, SL-0093:0094, SL-0101:0101, SL0151:0152, SL-0156:0158, SL-0205:0206, SL-0220, SL-0231:0232, SL-0327, SL-0524, SL-0535, SL-0174:0175, SL-0340; peroxisomes (113 proteins): SL-204; vesicles (748 proteins): SL-0498; plasma membrane (3173 proteins): SL-0039; cytoskeleton and cell adhesion (2161 proteins): SL-0009, SL-0090, SL-0092, SL-0118, SL-0280, SL-0501; cytoplasm (5319 proteins): SL-0086; secreted (1926 proteins): SL-0243-0245, SL-0347.

### Quantification of acetyl-CoA, succinyl-CoA, FAD, ATP, and ADP by LC-MS

Cultured cells were harvested by washing with ice cold PBS, dissociated with ice cold trypsin-EDTA (Gibco), and neutralized with ice cold culture media. Cells were pelleted at 700 × g at 4°C and washed with ice cold PBS. Cell pellets were then resuspended in 2.5% 5-sulfosalicylic acid (SSA) spiked with 10 µg of non-hydrolysable succinyl-CoA standard per sample. Samples were subjected to centrifugation at 17,000 × g at 4°C for 5 min, and supernatants were transferred into glass autosampler vials.

Samples were analyzed with a 1260 Infinity II HPLC coupled to an Agilent 6150 single quadrupole LC/MS. 10 µL of each sample was injected onto an Agilent InfinityLab Poroshell 120 EC-C18 column (699675–742). The auto-sampler temperature was set at 4°C and the column compartment temperature was set at 40°C. The mobile phase was composed of solvent A (50 mM formic acid in LCMS-grade H_2_O, adjusted to pH 8.2 with ammonium hydroxide) and solvent B (100% LCMS-grade methanol). The chromatographic gradient was run at a flow rate of 0.3 mL/min as follows: 0-1 min: hold at 20% B; 1–11 min: linear gradient from 20% to 100% B; 11–12 min: hold at 100% B; 12–13 min: linear gradient from 100% to 0% B; 13–20 min: hold at 0% B. The mass spectrometer was operated in selected ion monitoring (SIM), positive ion mode with the capillary voltage set to 3.5 kV, the nozzle voltage set to 2 kV. The sheath gas was held at 250°C at the flow rate 10 L/min and the drying gas was held at 300°C at the flow rate 5 L/min. The nebulizer pressure was set to 20 psi. The retention time and position of the peaks were confirmed using pure appropriate standard compounds freshly prepared in HPLC-grade water. The peak area/height was quantified using Agilent ChemStation. The mass to charge ratio (m/z) of the following ions were detected: acetyl-CoA (m/z 810.6), succinyl-CoA (m/z 868.6), FAD (m/z 786.5), ATP (m/z 508.2) and ADP (m/z 428.2).

### Quantification of NAD^+^ and NADH by LC-MS

Cultured cells were harvested by washing with ice cold PBS, dissociated with ice cold trypsin-EDTA (Gibco), and neutralized with ice cold culture media. Cells were pelleted at 700 × g at 4°C and washed with ice cold PBS. Cell pellets were then resuspended in 10% trichloroacetic acid (TCA) spiked with 10 µg ^13^C_5_-NAD^+^ standard per sample. Samples were subjected to centrifugation at 17,000 × g at 4°C for 5 min, and supernatants were transferred into new tubes. Metabolites were extracted using a 3:1 mixture of trichlorotrifluoroethane/trioctylamine. After vortex mixing and allowing phase separation to occur, the top layer was transferred to new tubes and adjusted to pH 8.0 with 1M Tris base before transferring into glass autosampler vials.

Samples were analyzed with a 1260 Infinity II HPLC coupled to an Agilent 6150 single quadrupole LC/MS. 10 µL of each sample was injected into an Agilent InfinityLab Poroshell 120 EC-C18 column (699675–742). The auto-sampler temperature was set at 4°C and the column compartment temperature was set at 40°C. The mobile phase was composed of solvent A (0.1% formic acid in LCMS-grade H_2_O) and solvent B (0.1% formic acid in LCMS-grade acetonitrile). The chromatographic gradient was run at a flow rate of 0.5 mL/min as follows: 0-2 min: hold at 0% B; 2–6 min: linear gradient from 0% to 60% B; 6–7 min: hold at 60% B; 7–9 min: linear gradient from 60% to 0% B; 9–19 min: hold at 0% B. The mass spectrometer was operated in selected ion monitoring (SIM), positive ion mode with the capillary voltage set to 3.5 kV, the nozzle voltage set to 2 kV. The sheath gas was held at 250°C at the flow rate 10 L/min and the drying gas was held at 300°C at the flow rate 5 L/min. The nebulizer pressure was set to 20 psi. The retention time and position of the peaks were confirmed using pure appropriate standard compounds freshly prepared in HPLC-grade water. Mass detection was completed in positive ESI mode. The peak area/height was quantified using Agilent ChemStation. The mass-to-charge ratio (m/z) of the following ions were detected: NAD^+^ (m/z 664.4), ^13^C_5_-NAD^+^ (m/z 669.4), NADH (m/z 666.4).

### FAO activity assessment by GC-MS

FAO activity was measured by monitoring the conversion rate of [U-[^13^C]]-palmitate (Cambridge Isotype Laboratories) to [M + 2]-labeled [^13^C] citrate as previously described (21, 69). Briefly, cultured cells were incubated with 100 µM [U-[^13^C]]-palmitate-BSA or unconjugated BSA control overnight. After incubation, cells were harvested by washing with ice cold PBS, dissociated with ice cold trypsin-EDTA (Gibco), and neutralized with ice cold culture media. Cells were pelleted by centrifugation at 700 × g at 4°C and washed with ice cold PBS. Cell pellets were lysed in 80% methanol and centrifuged to pellet debris. The resulting supernatants were dried with nitrogen gas, dissolved in 75 μL dimethylformamide, derivatized with 75 μL N-methyl-N-(tertbutyldimethylsilyl)trifluoroacetamide with 1% tertbutyldimetheylchlorosilane (Regis Technologies) and incubated at room temperature for 30 min. Samples were analyzed using an Agilent 7890B GC coupled to a 5977A mass detector.

Derivatized sample (3 µL) was injected into an Agilent HP-5ms Ultra Inert column, and the gas chromatograph oven temperature increased at 15°C/min up to 215°C, followed by 5°C/min up to 260°C, and finally at 25°C/min up to 325°C. The mass spectrometer was operated in splitless mode with electron impact mode at 70 eV. A mass range of 50–700 was analyzed, recorded at 1562 mass units/s. Data were analyzed using Agilent MassHunter Workstation Analysis. IsoPat2 software was used to adjust for natural abundance with unconjugated-BSA controls as previously performed (21). The extent of isotopic [^13^C] labeling in citrate and succinate was further divided by the percentage isotopic enrichment of intracellular [U-[^13^C]]-palmitate to determine the conversion rate of [U-[^13^C]]-palmitate to [M + 2]-labeled citrate or succinate in cells. Palmitate, citrate, and succinate were detected by gas chromatography/mass spectrometry as tert-butyldimetheylchlorosilane derivatives at the following mass-to-charge (m/z) values: endogenous palmitate (m/z 313), [U-[^13^C]]-palmitate (m/z 329), citrate (m/z 591– 593), and succinate (m/z 289-292).

### Statistical analysis

Details relating to all statistical analyses are described in the figure legends.

## Abbreviations

ATP: adenosine triphosphate
ADP: adenosine diphosphate
GTP: guanosine triphosphate
FAD(H_2_): flavin adenine dinucleotide
CoA: coenzyme A
imCC: immortalized mouse chromaffin cell
iMEF: immortalized mouse embryonic fibroblast
NAD(H): nicotinamide adenine dinucleotide
TCA: tricarboxylic acid
SDH: succinate dehydrogenase
FAO: fatty acid oxidation
BCAA: branched chain amino acid
PPGL: pheochromocytoma and paraganglioma
ETC: electron transport chain
DNA: deoxyribonucleic acid
PCA: principal component analysis
HIF: hypoxia-inducible factor
FDR: false discovery rate
HDAC: histone deacetylase
HAT: histone acetyltransferase
IPA: Ingenuity Pathway Analysis
mRNA: messenger ribonucleic acid
LCMS: liquid chromatography-mass spectrometry

## Declarations

## Ethics approval and consent to participate

Not applicable

## Consent for publication

Not applicable

## Availability of data and materials

The mass spectrometry proteomics data have been deposited to the ProteomeXchange Consortium (http://proteomecentral.proteomexchange.org) via the PRIDE partner repository (70) with the dataset identifier PXD066626 and DOI 10.6019/PXD066626. Materials and data are available upon request to the authors.

## Authors’ contributions

SXZ and LJM designed the experiments; SXZ, BM, MCM performed the experiments; SXZ, BM analyzed data; SXZ and LJM wrote the manuscript; TH and JF provided research materials and editing. All authors read and approved the final manuscript.

## Acknowledgments

We thank members of the Maher laboratory for their support. We acknowledge the assistance of the Mayo Clinic Flow Cytometry Core and of the Mayo Clinic Proteomics Core, which is a shared resource of the Mayo Clinic Cancer Center (NCI P30 CA15083). We also extend appreciation to Ya Li and Tianna Espe from the Hitosugi lab for technical assistance with metabolite quantification. We thank Peter Schultz (Scripps Research Institute) for the gift of pcDNA-mCherry-EGFP(K85TAG) and pCMVAcKRS-tRNAPyl plasmids used for sirtuin activity measurements. Some figure creation was facilitated by BioRender.com.

## Additional Materials

Additional file 1.pdf

PDF

Supplementary Material 1

Supplementary Figures S1-S8 and Supplementary Tables S1, S3, S4.

Additional file 2.xlsx

XLSX spreadsheet

Supplemental Table S2.

Top three clusters from DAVID functional annotation of disproportionately acylated proteins in imCCs and iMEFs.

The most significantly enriched (B-H corrected p-value < 0.05) pathways are shown for each cluster (up to five pathways if five or more pathways were included in a cluster).

Additional file 3.xlsx

XLSX spreadsheet

Supplemental Table S5.

Histone peptide acetylation and succinylation sites identified from imCCs and iMEFs.

