## Supplemental materials for "Distinct proteomic and acylproteomic adaptations to succinate dehydrogenase loss in two cell contexts"

### Table of Contents

#### Supplemental Figures

Supplemental Figure S1. Workflow for total and acylproteomic studies.
Supplemental Figure S2. TMT-based proteomic triplicates group by SDH status on principal component analysis (PCA) and for iMEFs are positively correlated with SILAC-based iMEF proteomics.
Supplemental Figure S3. Total proteomes and transcriptomes of imCCs and iMEFs with and without SDH loss are positively correlated.
Supplemental Figure S4. Ingenuity pathway analysis (IPA) shows enrichment of mitochondrial pathways in imCC total proteome and acetylome.
Supplemental Figure S5. SDH loss leads to downregulation of transcription-related, DNA damage, and replication-related pathways.
Supplemental Figure S6. SDH loss leads to increased fatty acid metabolism in imCCs but not in iMEFs.
Supplemental Figure S7. SDH loss leads to increased TCA cycle, Complex III, and Complex V expression in imCCs but not iMEFs.
Supplemental Figure S8. Protein acylation favors nuclear and cytoplasmic compartments over the secretory pathway in imCCs and iMEFs.
Supplemental Figure S9. SDH loss leads to upregulation and hyperacylation of TCA cycle and ETC proteins in imCCs but not in iMEFs.

#### Supplemental Tables

Supplemental Table S1. IPA pathways upregulated in imCCs and downregulated in iMEFs upon SDH loss.
Supplemental Table S2. Top three clusters from DAVID functional annotation of disproportionately acylated proteins in imCCs and iMEFs.
Supplemental Table S3. Histone acetyltransferases (HATs) are predominantly downregulated in imCCs but not in iMEFs upon SDH loss.
Supplemental Table S4. Histones identified in imCCs and iMEFs.
Supplemental Table S5. Histone peptide acetylation and succinylation sites identified in imCCs and iMEFs.

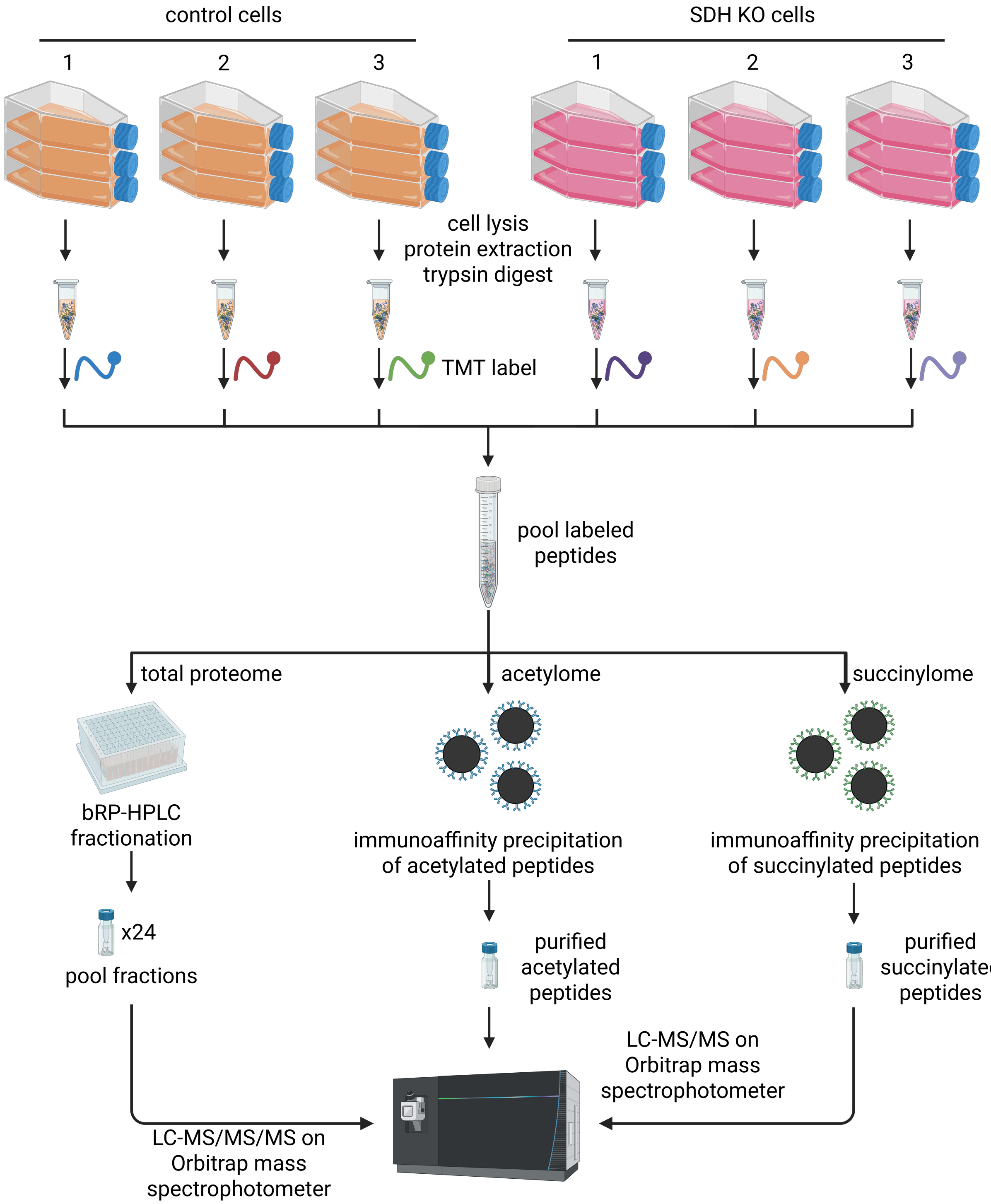


**Supplemental Figure S1. Workflow for total and acylproteomic studies.**

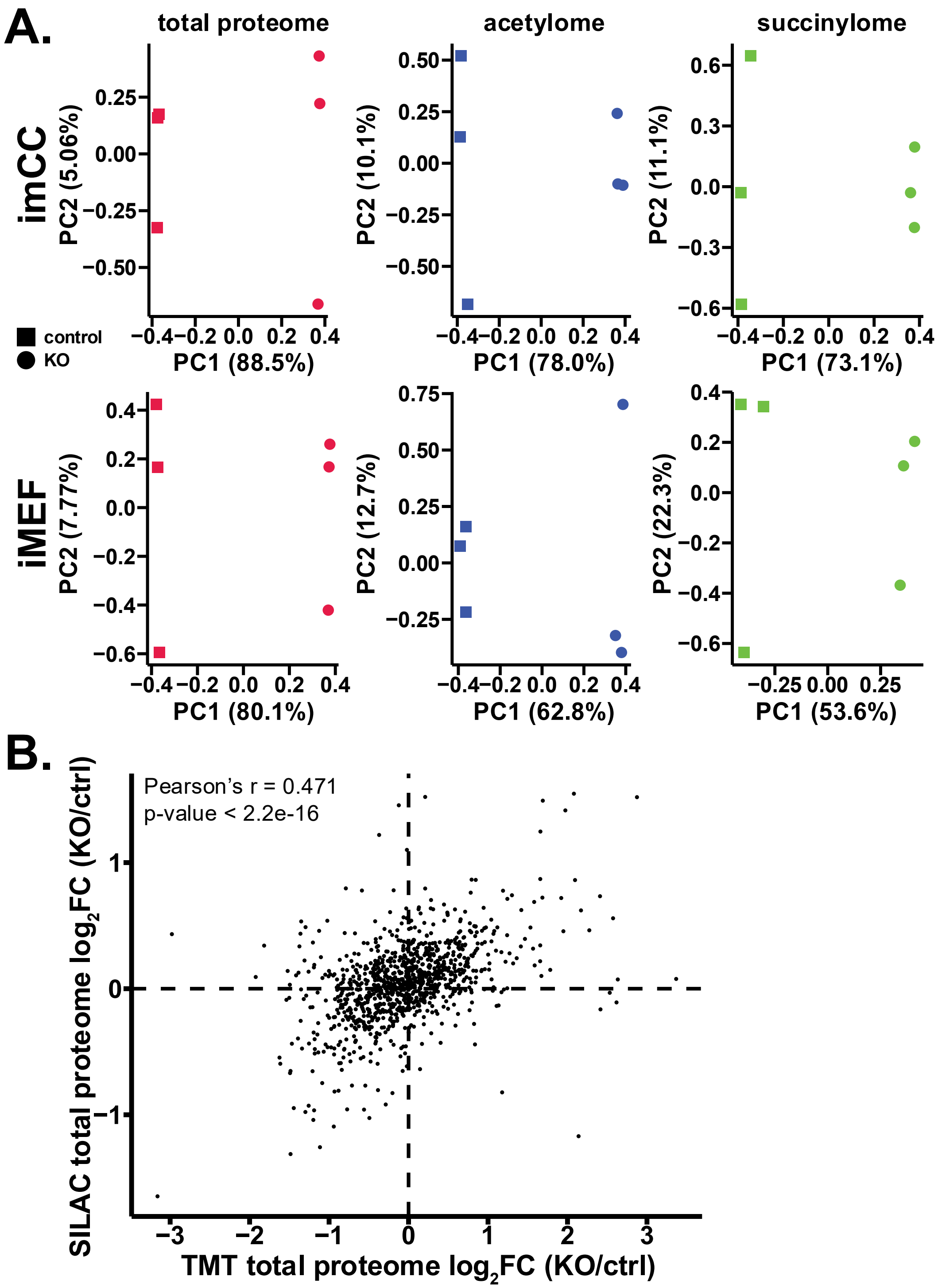


**Supplemental Figure S2. TMT-based proteomic triplicates group by SDH status on principal component analysis (PCA) and for iMEFs are positively correlated with SILAC-based iMEF proteomics. A.** PCA of total proteome, acetylome, and succinylome in imCCs and iMEFs. **B.** Correlation plot of changes upon SDH loss in iMEF total proteome from TMT-based proteomics to previous iMEF total proteome from SILAC-based proteomics (21).

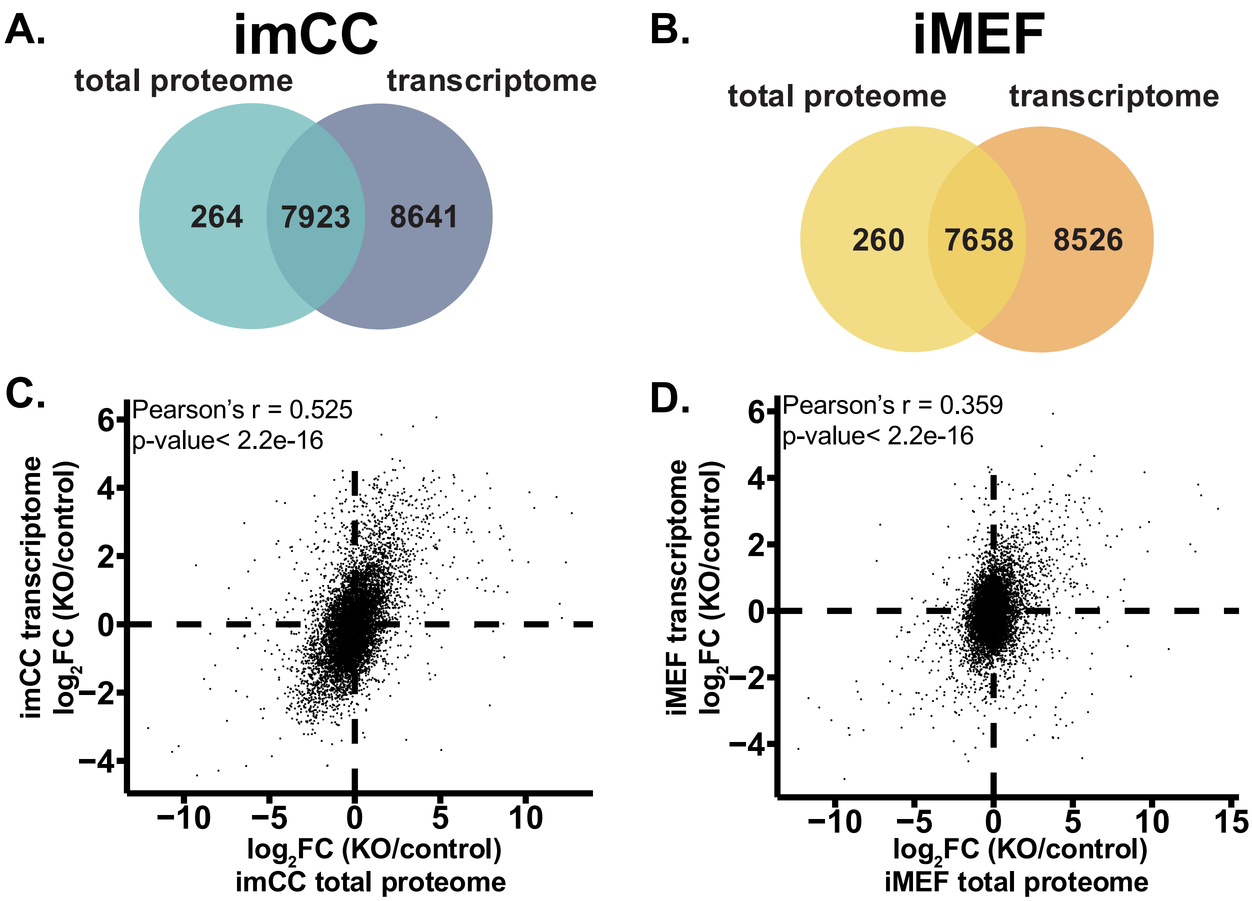


**Supplemental Figure S3. Total proteomes and transcriptomes of imCCs and iMEFs with and without SDH loss are positively correlated. A-B.** Venn diagram of protein-mRNA pairs between **A.** imCC and **B.** iMEF total proteome and transcriptome. **C-D.** Correlation plot between changes in total proteome and transcriptome upon SDH loss in **C.** imCCs and **D.** iMEFs. Transcriptomes from (37).

A.

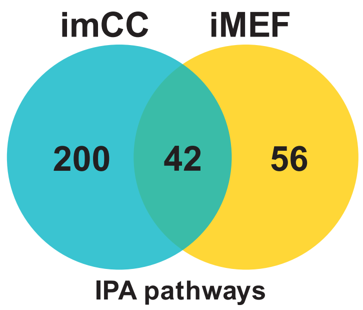


B.

**Regulation of Lipid Metabolism by PPARα**

| | imCC | | iMEF | |
|---|---|---|---|---|
| | z-score | $-\log_{10}$(B-H corrected p-value) | z-score | $-\log_{10}$(B-H corrected p-value) |
| | -2.2 | 3.3 | 2.138 | 2.31 |
| proteins | ACADM, ACSL1, ANKRD1, CARM1, CPT1A, FHL2, HMGCR, HMGCS1, MED10, MED11, MED15, MED17, MED19, MED22, MED27, MED28, MED29, MED30, MED31, MED4, MED9, RGL1, SLC27A1, SREBF1, TGS1 | | ACOX1, ACSL1, ANGPTL4, ARNT2, CDK19, FDFT1, FHL2, GLIPR1, HMGCR, HMGCS1, MED18, NCOA1, NCOA2, RGL1 | |

C.

| | imCC | |
|---|---|---|
| **IPA Canonical Pathway** | z-score | $-\log_{10}$(B-H corrected p-value) |
| Mitochondrial iron-sulfur cluster biogenesis | 2 | 1.41 |
| Maturation of TCA enzymes and regulation of TCA cycle | 2.121 | 3.26 |
| Mitochondrial Division Signaling Pathway | 2.611 | 3.67 |
| Mitochondrial protein degradation | 3.578 | 2.6 |
| Mitochondrial Fatty Acid Beta-Oxidation | 3.742 | 4.98 |

D. E. F. G.

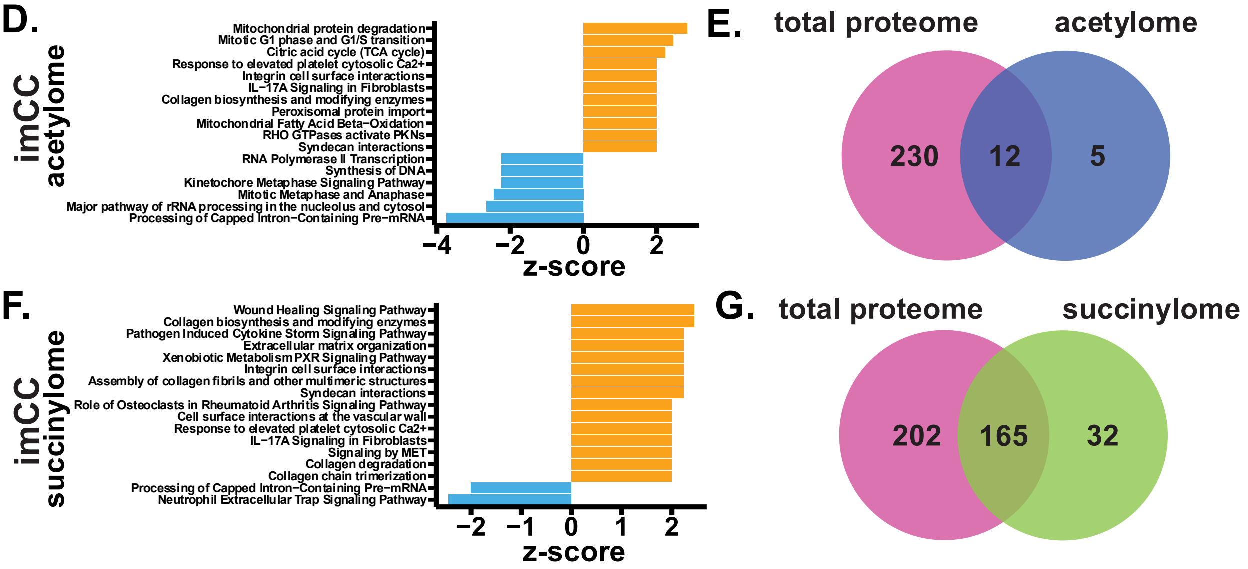


**Supplemental Figure S4. Ingenuity pathway analysis (IPA) shows enrichment of mitochondrial pathways in imCC total proteome and acetylome. A.** Venn diagram of IPA pathways altered by SDH loss, shared and unique to imCC and iMEF total proteomes. **B.** IPA pathway scores and protein hits for “Regulation of Lipid Metabolism by PPARα” for imCC and iMEF total proteomes. Mediator subunits are shown in red. **C.** Mitochondrial IPA pathways present in imCC total proteome dysregulated pathways. **D-E.** Top 15 upregulated and downregulated pathways for **D.** acetylome and **E.** succinylome in imCCs. **F-G.** Comparison of shared and unique pathways between total proteome and **F.** acetylome or **G.** succinylome pathways for imCCs.

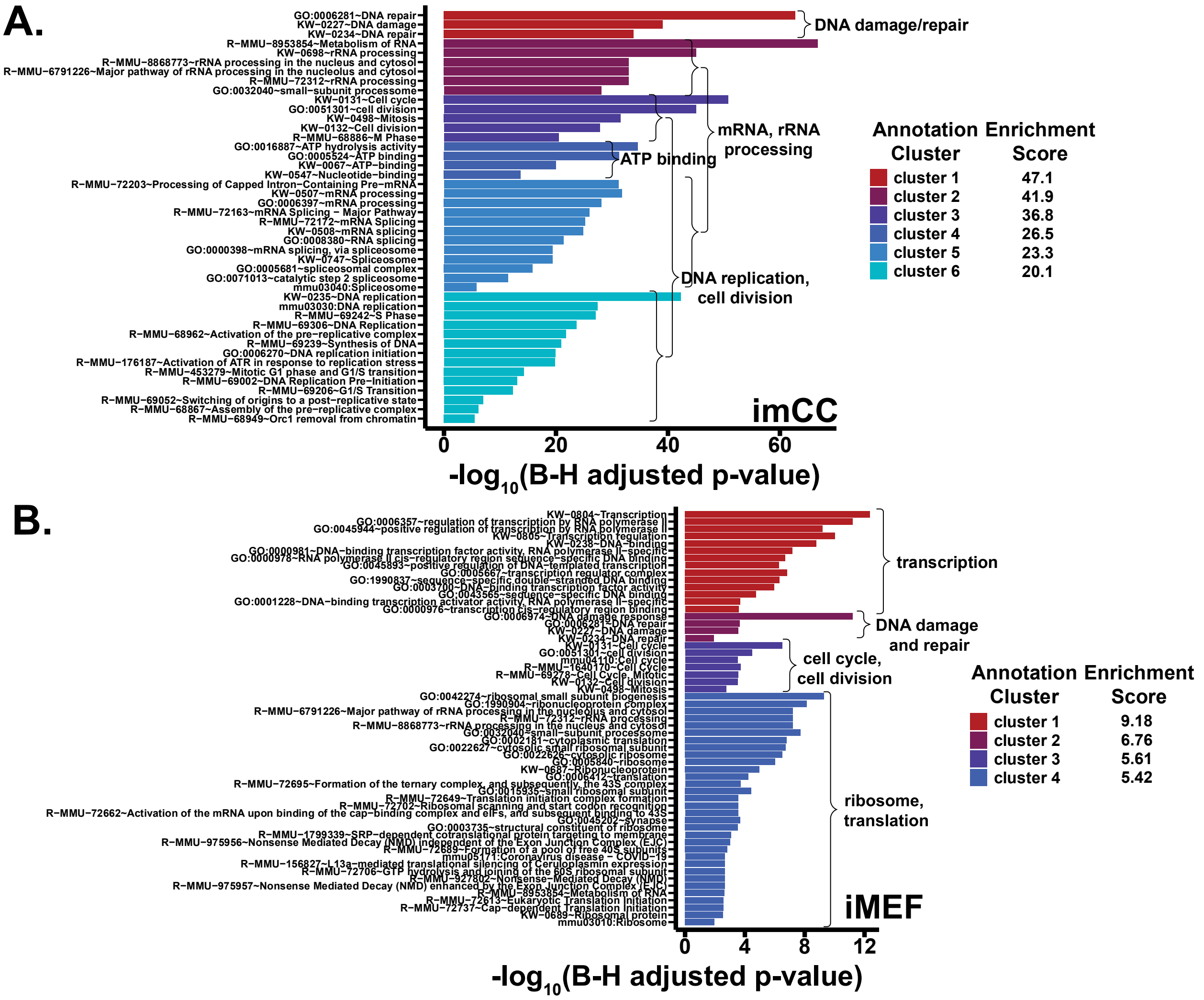


**Supplemental Figure S5. SDH loss leads to downregulation of transcription-related, DNA damage, and replication-related pathways. A-B.** Top six annotation clusters from DAVID Functional Analysis of nuclear proteins with FC < -2, p-value < 0.05 in **A.** imCCs and **B.** iMEFs.

**A.**

| Protein | Accession Number | imCC Fold Change (KO/control) | imCC FDR-adjusted p-value | iMEF Fold Change (KO/control) | iMEF FDR-adjusted p-value |
|---|---|---|---|---|---|
| CPT1A | P97742 | 5.15 | 1.2E-05 | -1.07 | 0.26 |
| PPARD | P35396 | N/A | N/A | N/A | N/A |
| RXRA | P28700 | 1.07 | 0.003 | 1.54 | 0.00023 |
| SLC25A20 | Q9Z2Z6 | 1.80 | 0.00034 | 1.27 | 0.039 |
| ACACA | Q5SWU9 | -2.62 | 2.8E-05 | -1.30 | 0.00043 |
| THRSP | Q62264 | N/A | N/A | N/A | N/A |
| MID1IP1 | Q9CQ20 | -1.98 | 9.5E-05 | -1.32 | 0.008 |
| SLC22A5 | Q9Z0E8 | N/A | N/A | N/A | N/A |
| CPT1B | Q924X2 | N/A | N/A | N/A | N/A |
| ACACB | E9Q4Z2 | -1.88 | 0.00011 | N/A | N/A |
| CPT2 | P52825 | 1.82 | 0.00004 | 1.78 | 0.00029 |
| PRKAA2 | Q8BRK8 | N/A | N/A | -1.05 | 0.42 |
| PRKAB2 | Q6PAM0 | 1.99 | 0.00024 | 1.23 | 0.020 |
| PRKAG2 | Q91WG5 | 1.25 | 0.0035 | N/A | N/A |
| ACBD6 | Q9D061 | -1.99 | 2.2E-05 | 1.15 | 0.022 |
| MCAT | Q8R3F5 | 2.87 | 3.8E-06 | 1.43 | 0.00 |
| NDUFAB1 | Q9CR21 | 1.35 | 0.022 | N/A | N/A |
| ACSF2 | Q8VCW8 | 2.23 | 4.7E-06 | 1.51 | 0.00013 |
| MECR | Q9DCS3 | 1.48 | 6.6E-05 | -1.67 | 0.00036 |
| ACADM | P45952 | 3.06 | 4.5E-06 | -1.20 | 0.0066 |
| ECHS1 | Q8BH95 | 6.25 | 2.3E-06 | -1.07 | 0.061 |
| HADH | Q61425 | 2.66 | 3.6E-06 | -1.04 | 0.39 |
| HADHA | Q8BMS1 | 1.48 | 1.5E-05 | 1.57 | 0.0017 |
| HADHB | Q99JY0 | 1.51 | 0.00072 | 1.24 | 0.0074 |
| ACADL | P51174 | 2.22 | 8.1E-06 | 1.56 | 0.00025 |
| ACADVL | P50544 | 1.39 | 5.2E-05 | 1.40 | 0.00064 |
| ACADS | Q07417 | 1.63 | 6.2E-05 | 2.67 | 2.9E-05 |
| PCCB | Q99MN9 | 2.22 | 1.4E-05 | 1.26 | 0.0024 |
| PCCA | Q91ZA3 | 1.69 | 6.7E-05 | -1.08 | 0.17 |
| MCEE | Q9D1I5 | 4.54 | 4.9E-05 | N/A | N/A |
| MMUT | P16332 | 2.66 | 1.6E-05 | 1.13 | 0.10 |
| MMAA | Q8C7H1 | 2.45 | 5.4E-06 | 1.52 | 0.022 |
| ACAA2 | Q8BWT1 | 2.60 | 1.3E-05 | 1.15 | 0.026 |
| ACAD10 | Q8K370 | 7.54 | 2.3E-06 | 7.58 | 4.5E-05 |
| THEM4 | Q3UUI3 | 1.44 | 0.00086 | N/A | N/A |
| ACOT2 | Q9QYR9 | 19.46 | 3.2E-06 | 3.14 | 0.00003 |
| ACOT1 | O55137 | 4.60 | 4.7E-05 | -1.26 | 0.017 |
| ACOT5 | Q6Q2Z6 | N/A | N/A | N/A | N/A |
| ACOT3 | Q9QYR7 | N/A | N/A | N/A | N/A |
| ACOT9 | Q9R0X4 | -1.41 | 4.1E-05 | -1.11 | 0.10 |
| THEM5 | Q9CQJ0 | N/A | N/A | N/A | N/A |
| ACBD7 | Q9D258 | N/A | N/A | N/A | N/A |
| DBI | P31786 | 1.45 | 0.049 | 2.25 | 0.0024 |
| ACOT7 | Q91V12 | -1.29 | 0.001 | -2.48 | 3.3E-05 |
| ACOT12 | Q9DBK0 | N/A | N/A | N/A | N/A |
| ACOT13 | Q9CQR4 | -1.02 | 0.186 | 1.92 | 8.2E-05 |
| ACOT11 | Q8VHQ9 | N/A | N/A | N/A | N/A |
| PCTP | P53808 | 1.22 | 0.0017 | N/A | N/A |
| ECI1 | P42125 | 2.11 | 7.9E-06 | 1.76 | 0.0011 |
| DECR1 | Q9CQ62 | 2.17 | 3.5E-05 | -1.09 | 0.028 |
| ACAD11 | Q80XL6 | -1.12 | 0.034 | -1.05 | 0.19 |

**B.**

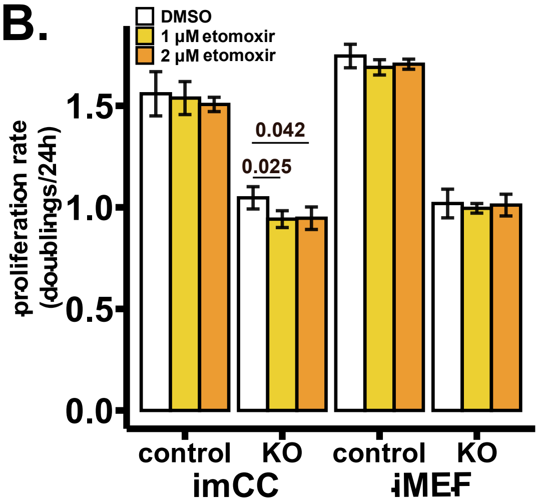


**Supplemental Figure S6. SDH loss leads to increased fatty acid metabolism in imCCs but not in iMEFs. A.** Changes in carnitine shuttle (R-MMU-200425) and FAO (R-MMU-77289) protein levels upon SDH loss from total proteomics. **B.** Proliferation rates of vehicle (DMSO) and etomoxir-treated imCCs and iMEFs in standard culturing media. Cells were treated at indicated concentrations for 3 cell doublings. P-values were calculated using two-tailed Welch's t-tests. Data representative of 3 independent replicates.

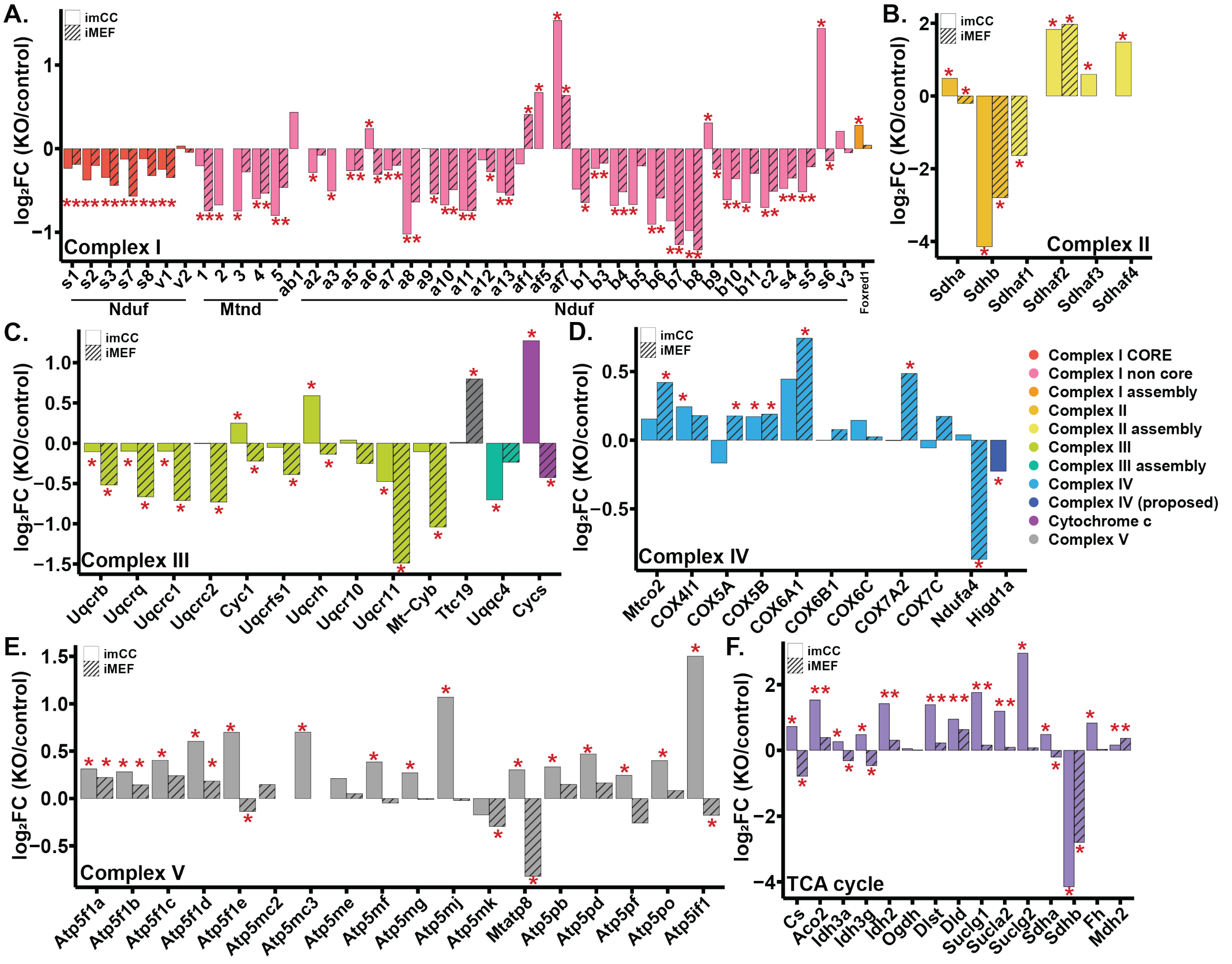


**Supplemental Figure S7. SDH loss leads to increased TCA cycle, Complex III, and Complex V expression in imCCs but not in iMEFs. A-E.** Fold change in expression of electron transport chain complexes and assembly factors for **A.** Complex I, **B.** Complex II, **C.** Complex III and cytochrome c, **D.** Complex IV, and **E.** Complex V upon SDH loss in imCCs and iMEFs. **F.** Fold change (FC) in TCA cycle protein expression upon SDH loss in imCCs and iMEFs. Dashed lines indicate |FC| = 1. Red asterisks indicate P (FDR-adjusted p-value) < 0.05.

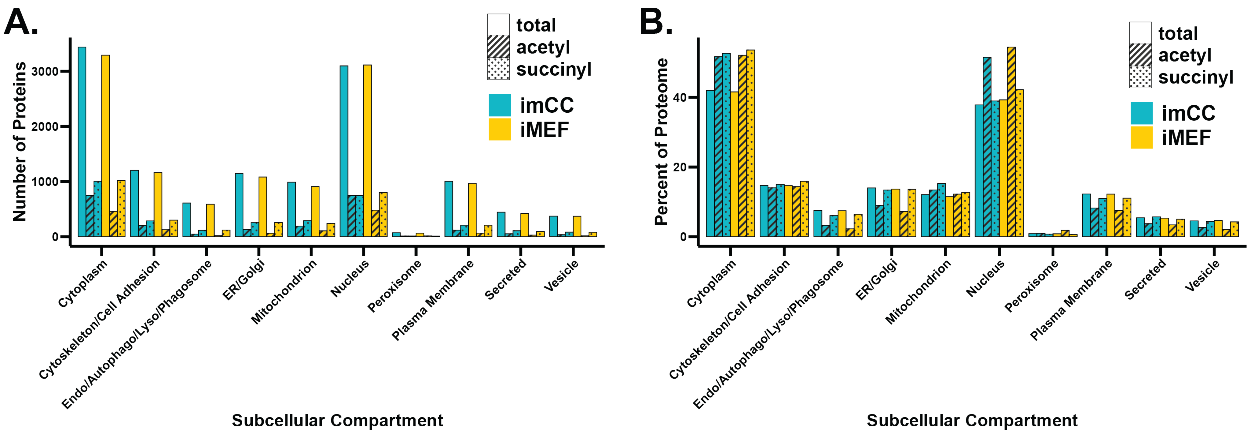


**Supplemental Figure S8. Protein acylation favors nuclear and cytoplasmic compartments over the secretory pathway in imCCs and iMEFs. A-B.** Distribution of proteins detected in proteomic studies by primary subcellular compartment by **A.** total number and **B.** percentage of corresponding quantified proteome. Only proteins with annotated locations in the UniProt keyword database are included.

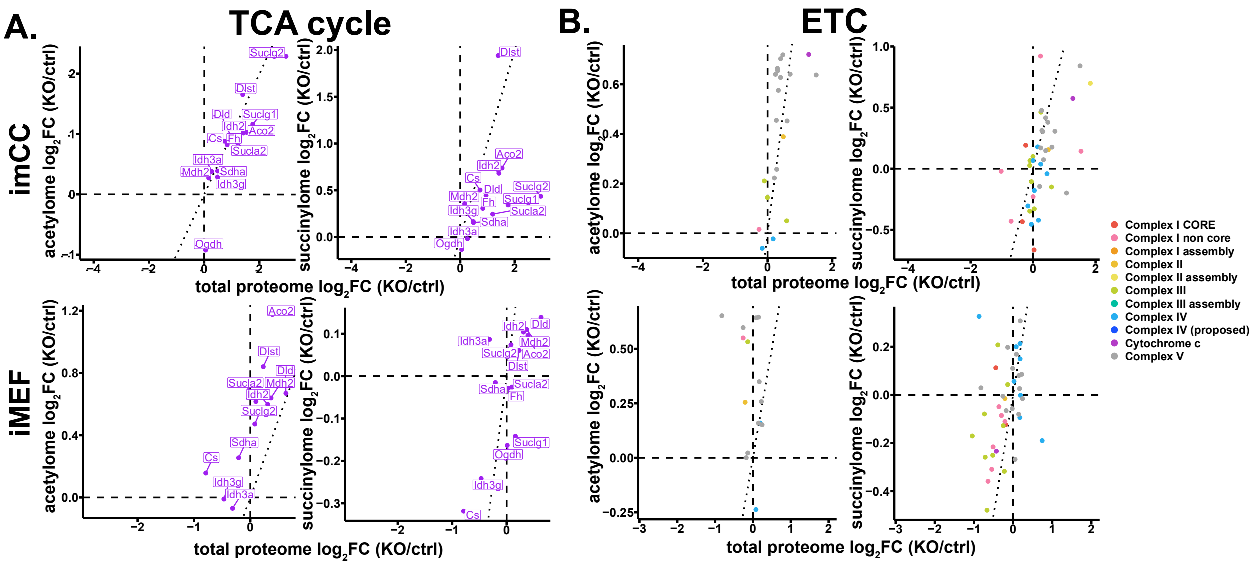


**Supplemental Figure S9. SDH loss leads to upregulation and hyperacylation of TCA cycle and ETC proteins in imCCs but not in iMEFs. A-B.** Correlation plots comparing changes in total and acylated proteins in imCCs and iMEFs in **A.** TCA cycle proteins, and **B.** ETC proteins. Dotted diagonal line shows y=x where there is a 1:1 correlation of total proteome and acylome. Dotted diagonal line shows y=x where there is a 1:1 correlation of total proteome and acylome.

**Supplemental Table S1. IPA pathways upregulated in imCCs and downregulated in iMEFs upon SDH loss.**

| | imCC | | iMEF | |
|---|---|---|---|---|
| **IPA Canonical Pathway** | **z-score** | **$-log_{10}$(B-H corrected p-value)** | **z-score** | **$-log_{10}$(B-H corrected p-value)** |
| Signaling by Rho Family GTPases | 2.082 | 6.55 | -2.524 | 1.99 |
| VEGF Family Ligand-Receptor Interactions | 2.138 | 1.49 | -2.333 | 1.76 |
| Ovarian Cancer Signaling | 2.324 | 6.54 | -2.496 | 6.95 |
| Platelet homeostasis | 2.4 | 3.79 | -2.111 | 2.19 |
| Paxillin Signaling | 2.558 | 7.16 | -2.333 | 1.93 |
| HIF1α Signaling | 2.596 | 4.22 | -3.128 | 3.36 |
| Molecular Mechanisms of Cancer | 2.67 | 2.07 | -3.15 | 2.22 |
| IL-8 Signaling | 2.722 | 5.78 | -3.273 | 2.69 |
| RHOA Signaling | 2.746 | 4.25 | -2.714 | 1.47 |
| Hereditary Breast Cancer Signaling | 2.777 | 11.2 | -2.828 | 3.34 |
| Signaling by VEGF | 2.837 | 2.49 | -2.668 | 4.23 |
| Eicosanoid Signaling | 2.949 | 3.32 | -2.921 | 4.21 |
| GNRH Signaling | 2.959 | 4.39 | -3.153 | 2.4 |
| VEGF Signaling | 3.13 | 4.83 | -2.53 | 1.81 |
| Extra-nuclear estrogen signaling | 3.153 | 2.8 | -3 | 1.71 |
| Ephrin Receptor Signaling | 3.157 | 6.15 | -2.496 | 4.89 |

**Supplemental Table S2. Top three clusters from DAVID functional annotation of disproportionately acylated proteins in imCCs and in iMEFs.**
The most significantly enriched (B-H corrected p-value < 0.05) pathways are shown for each cluster (up to five pathways if five or more pathways were included in a cluster).

See separate Excel file.

**Supplemental Table S3. Histone acetyltransferases (HATs) are predominantly downregulated in imCCs but not in iMEFs upon SDH loss.**

| | | | imCC | | iMEF | |
|---|---|---|---|---|---|---|
| **HAT family** | **HAT** | **Accession Numbers** | **Fold Change (KO/control)** | **FDR-adjusted p-value** | **Fold Change (KO/control)** | **FDR-adjusted p-value** |
| GNAT | KAT2A | Q9JHD2 | -2.45 | 6.35E10-6 | 2.44 | 0.017 |
| | KAT2B | Q9JHD1 | N/A | N/A | 2.09 | 0.0081 |
| | HAT1 | Q8BY71 | -5.71 | 2.63E10-6 | -1.62 | 0.00036 |
| MYST | KAT5 | Q8CHK4 | -2.10 | 0.00030 | 1.36 | 0.007 |
| | KAT6A | Q8BZ21 | -1.25 | 0.0042 | 1.03 | 0.25 |
| | KAT7 | Q5SVQ0 | -2.05 | 1.62E10-5 | -1.48 | 0.0029 |
| | KAT8 | Q9D1P2 | -1.46 | 9.14E10-5 | 1.11 | 0.073 |
| p300 | p300 | B2RWS6 | -1.26 | 0.00030 | 1.07 | 0.028 |
| | CREBBP | P45481 | -1.39 | 3.72E10-5 | 1.31 | 2.44E10-5 |
| Other | ATF2 | P16951 | -1.48 | 0.00032 | 1.25 | 0.12 |
| | CLOCK | O08785 | -1.43 | 0.0017 | 1.06 | 0.63 |

**Supplemental Table S4. Histones identified in imCCs and iMEFs.**

| Histone | Accession Numbers | imCC | iMEF |
|---|---|---|---|
| H1.0 | P10922 | yes | yes |
| H1.1 | P43275 | yes | yes |
| H1.2 | P15864 | yes | yes |
| H1.3 | P43277 | yes | yes |
| H1.4 | P43274 | yes | yes |
| H1.5 | P43276 | yes | yes |
| H2A type 1-F | Q8CGP5 | no | yes |
| H2A type 1-G | C0HKE5 | yes | no |
| H2A type 2-A | Q6GSS7 | yes | no |
| H2A type 2-B | Q64522 | yes | yes |
| H2A type 2-C | Q64523 | no | yes |
| H2A.J | Q8R1M2 | yes | yes |
| H2A.V | Q3THW5 | yes | no |
| H2A.X | P27661 | yes | yes |
| H2A.Z | P0C0S6 | yes | yes |
| H2B type 1-B | Q64475 | yes | yes |
| H2B type 1-K | Q8CGP1 | yes | yes |
| H2B type 2-B | Q64525 | yes | yes |
| H2B type 2-E | Q64524 | yes | no |
| H2B type 3-B | Q8CGP0 | yes | no |
| H2B.U | Q9D2U9 | yes | yes |
| H3.1 | P68433 | yes | yes |
| H3.2 | P84228 | yes | yes |
| H3.3 | P84244 | yes | yes |
| H3.3C | P02301 | yes | yes |
| H4 | P62806 | yes | yes |

**Supplemental Table S5. Histone peptide acetylation and succinylation sites identified from imCCs and iMEFs.**

See separate Excel file.
